# The miR-615-5p/ID1 Axis Regulates Autophagy-Associated Migration in Pancreatic Ductal Adenocarcinoma

**DOI:** 10.64898/2026.08.27.747461

**Authors:** Abhishek Sarkar, Subha Ray, Aishwarya Ray, Kaushik Biswas

## Abstract

Pancreatic ductal adenocarcinoma (PDAC) is an aggressive malignancy characterized by high metastatic dissemination, therapy resistance, and poor clinical outcome. Inhibitor of differentiation 1 or ID1, is frequently overexpressed in PDAC and is associated with tumour progression and adverse clinical outcome. However, the mechanisms governing its post-transcriptional regulation remain insufficiently characterized. Here, we identify tumour-suppressive miR-615-5p as a regulator of ID1 expression in PDAC. Integrative *in-silico* target prediction prioritized miR-615-5p based on seed complementarity and thermodynamic stability with the ID1 3′-UTR. Expression analysis of available PDAC clinical datasets revealed reduced miR-615-5p expression associated with increased ID1 expression. Direct association was validated using luciferase reporter assays, where miR-615-5p suppressed 3’-UTR reporter activity of ID1 in a sequence dependent manner, while mutation of the predicted binding site attenuated this effect. Further biotinylated-RIP and AGO2-RIP assays demonstrated the co-enrichment of ID1 transcripts and miR-615-5p with AGO2 associated RISC complexes, while AntimiR mediated inhibition of miR-615-5p perturbs association between miR/ID1 to AGO2, supporting interaction specificity. Functionally, modulation of miR-615-5p altered ID1 expression and impacted PDAC cell migration *in vitro*. Mechanistic analyses further indicated that the miR-615-5p/ID1 axis influences autophagic flux where miR-615-5p mediated inhibition of autophagy suppresses ID1 dependent cellular migration. Collectively, these findings define a previously uncharacterized miRNA-dependent regulation of ID1 expression and link this axis to autophagy-associated migratory responses in PDAC cells. The study expands the post-transcriptional regulatory landscape of ID1 and provides a possible mechanism where suppression of miR-615-5p leads to ID1 overexpression and subsequent poor clinical outcome in PDAC cells.

**Statement of implication:** miR-615-5p negatively regulate ID1 expression by targeting its 3’-UTR and cellular migration by suppressing ID1 dependent autophagy in PDAC.

## Introduction

Pancreatic ductal adenocarcinoma (PDAC) arises from the ductal epithelium of the pancreas and accounts for more than 90% of pancreatic malignancies worldwide (1). It remains one of the most lethal solid tumours, characterized by late clinical presentation, rapid metastatic progression, and limited responsiveness to current therapeutic regimens, resulting in extremely poor survival outcomes (2–4). The aggressive clinical behaviour of PDAC reflects coordinated alterations in oncogenic signalling networks and stress-adaptation pathways, underscoring the need to identify regulatory mechanisms that control key tumour-promoting factors.

Inhibitor of differentiation 1 (ID1) belongs to the basic helix-loop-helix family of proteins (5) that modulate multiple cellular processes that are crucial for maintaining cellular homeostasis, including cellular proliferation, cell migration, angiogenesis, stem cell niche maintenance etc (5–8). ID1 overexpression has been reported across multiple malignancies, including PDAC, where elevated ID1 expression correlates with tumour aggressiveness and poor prognosis (6, 9, 10). ID1 promotes tumorigenesis by inducing pro-proliferative and pro-migratory activity while facilitating angiogenesis and contributing to therapy resistance. Considering its consistent overexpression and multi-dimensional role, in a recent study Tang et al. suggested ID1 as a potential malignant biomarker in PDAC (11).

As opposed to its function, the regulation of ID1 is complex and poorly understood. Multiple upstream signalling pathways, including TGF-β/Smad, PI3K-Akt, MEK-ERK, and BMP signalling, were reported to regulate ID1 transcription in various cancer types including PDAC (10, 12–14). In contrast, post-transcriptional regulatory mechanisms governing ID1 expression remains elusive. MicroRNAs (miRNAs) are small non-coding RNAs that mediate post-transcriptional gene regulation through mRNA destabilization or translational repression and frequently function as critical modulators of oncogenic networks(15, 16). Despite their widespread regulatory roles in cancer, evidence indicating miRNAs as potential regulators of ID1 post-transcriptional regulation is limited. So far, only two miRNAs, miR-29b and miR-381 were experimentally validated as direct regulators of ID1 through 3′-UTR targeting, in lung adenocarcinoma models (17–19). Hence, a comprehensive analysis of miRNAs regulating ID1 is crucial since ID1 acts as a key orchestrator of tumour progression, metastasis, and cancer stem cell (CSC) maintenance across various malignancies.

In this study, we considered ID1’s broad oncogenic relevance across multiple cancer types, where elevated ID1 expression is consistently associated with aggressive tumour behaviour and adverse clinical outcome. Despite its established oncogenic roles in other malignancies, ID1 particularly in PDAC is closely associated with metastatic progression and adaptive stress responses that support tumour persistence in a nutrient-limited microenvironment(11, 20). This makes ID1 an important regulatory node in PDAC biology and a compelling candidate for detailed investigation of its regulatory mechanism. To identify post-transcriptional regulators of ID1, a systematic *in silico* screening strategy followed by multiple orthogonal interaction and functional validation assays identified and established miR-615-5p as a regulator of ID1 gene expression. miR-615-5p was previously identified as a tumour-suppressive miRNA in PDAC and other cancer types, where it was shown to inhibit cellular migration, invasion, and tumour growth (21–24). Suppression of miR-615-5p expression in PDAC was attributed to promoter hypermethylation and concomitant epigenetic silencing. Prior functional studies demonstrated that miR-615-5p targets oncogenic effectors IGF2 resulting in suppressed cellular migration and proliferation, supporting its role as tumour suppressor in PDAC (21). These observations make miR-615-5p a biologically plausible upstream regulator of ID1.

PDAC tumours are further distinguished by elevated basal autophagy, which supports survival under conditions of metabolic stress and nutrient limitation characteristic of the pancreatic tumour microenvironment (25, 26). Autophagy enables metabolic rewiring and sustains tumour cell fitness by recycling intracellular components and maintaining energy balance(27). A recent study on ovarian cancer established a novel role of ID1 in promoting cellular autophagy to adapt therapeutic stress against cisplatin and paclitaxel, thereby potentially contributing to generation of resistance (28). This finding suggests a potential intersection between ID1 regulation and autophagy in aggressive tumours such as PDAC.

In the present study, *in silico* prediction followed by experimental validation, identified miR-615-5p as a negative regulator of ID1. Using reporter assays, miRNA pulldown, AGO2-RIP approaches, a sequence-specific association between miR-615-5p and the ID1 3′-UTR was validated. Functional analyses further indicate that miR-615-5p mediated ID1 suppression is associated with altered autophagic activity and reduced migratory potential in cellular models. The outcomes collectively demonstrate a novel regulatory mechanism, where the tumour suppressor miR-615-5p targets oncogenic ID1, thereby identifying a novel miR-615-5p/ID1 regulatory axis. Suppression of miR-615-5p expression leads to upregulated ID1 expression in PDAC, thereby promoting tumorigenic progression. This observation is consistent with the reported association between elevated ID1 expression and poor clinical outcomes in PDAC (1, 4). Defining the miR-615-5p–ID1 regulatory relationship advances the mechanistic understanding of post-transcriptional control of ID1 expression in PDAC. Considering the established association of ID1 with aggressive disease behaviour, elucidating its upstream miRNA regulation provides a mechanistic framework for further biological and translational evaluation. While translational implications require substantial additional validation, these results may provide a mechanistic basis for future studies evaluating the regulatory control of ID1 in PDAC.

## Materials and methods

### Analysis of ID1 gene expression in different cancers using GEPIA2.0

Gene Expression Profiling Interactive Analysis, version 2 (GEPIA2.0) (29) was utilized to analyse the gene expression profiles of ID1 between the tumour tissues and the corresponding normal tissues. The ‘profile’ module from the GEPIA2.0 database was used to carry out the analysis based on TCGA and Genotype-Tissue Expression dataset (GTEx) samples. The screening criteria in GEPIA2.0 were as follows: p < 0.05 and the cutoff of |Log2FC| was 1. Both the normal samples and the tumour samples were analysed using the one-way ANOVA.

### Prognosis and Survival analysis of ID1 in PAAD and PDAC using KMplotter

The prognostic importance of ID1 expression in PAAD and PDAC was assessed using the Kaplan-Meier Plotter (KMplotter) (30), an online tool (http://kmplot.com) that combines gene expression data and survival information from various datasets, including GEO, EGA and TCGA. Gene chip module was used to evaluate the association between ID1 expression with overall survival (OS) and disease free survival (DFS) in PAAD patients. To analyse the role of ID1 expression on the overall survival of PDAC, RNA-seq module was used. The dataset was filtered using the default settings unless stated otherwise. Patients were automatically divided into high and low expression groups based on the median expression value of the queried gene. Survival curves were created using the Kaplan-Meier method. The web-tool calculated and displayed the hazard ratio (HR) and log-rank p-value. Hazard ratios (HRs) with 95% confidence intervals (CIs) were calculated by KMplotter to assess the relative hazard of death between the high– and low-expression groups. An HR >1 indicated an increased hazard of death in the high-ID1 expressing group. Hazard ratios (HRs) were interpreted as the relative change in hazard, with the percentage change in hazard calculated as 100 × (HR − 1) (31). A p-value of less than 0.05 was considered statistically significant.

### Expression analysis of ID1 in PDAC and normal Pancreas using UCSC XENA

To compare ID1 expression between pancreatic ductal adenocarcinoma (PDAC) and normal pancreatic tissues, publicly available transcriptomic datasets were analysed using the UCSC Xena platform (32). Gene expression data were obtained from the integrated TCGA-TARGET-GTEx cohort, which provides uniformly processed RNA-seq datasets across tumour and normal tissues derived from The Cancer Genome Atlas (TCGA), Therapeutically Applicable Research to Generate Effective Treatments (TARGET), and Genotype-Tissue Expression (GTEx), enabling cross-cohort comparisons between malignant and non-malignant samples. Samples within the TCGA-TARGET-GTEx cohort were filtered using the metadata fields “study,” “primary site,” and “primary disease or tissue” to isolate pancreatic adenocarcinoma (PAAD) and normal pancreas datasets for downstream analysis. ID1 expression values were retrieved for 167 normal pancreatic samples from the GTEx cohort and 183 PAAD tumour samples from the TCGA cohort. To specifically define PDAC cases within the PAAD dataset, the TCGA-PAAD cohort was further stratified based on the “histological type” clinical annotation. Using this criterion, 150 samples classified as PDAC were identified. These PDAC cases were then cross-matched with the previously extracted PAAD tumour samples from the TCGA-TARGET-GTEx cohort to ensure consistency between clinical annotation and expression datasets. For all selected samples, log₂-transformed RSEM-normalized expression values [log₂(RSEM + 1)] for ID1 were obtained. For graphical representation, values were transformed to linear scale where appropriate and plotted to compare expression levels between normal pancreas and PDAC groups using GraphPad Prism software.

### Prediction of ID1 targeting microRNAs

To identify potential microRNAs (miRNAs) targeting the ID1 gene, *in silico* analysis was performed using three different miRNA target prediction tools: TargetScan (http://www.targetscan.org) (33), miRWalk (http://mirwalk.umm.uni-heidelberg.de/) (34), and miRSystem (http://mirsystem.cgm.ntu.edu.tw/) (35). Default parameters were applied to retrieve the predicted miRNAs based on seed region complementarity, evolutionary conservation, and binding site accessibility etc. The lists containing the predicted miRNAs that target ID1 were obtained from each database. To minimize the false positive results, the intersection set was manually curated to identify commonly predicted miRNAs by all three platforms. The depicted venn diagram was created using jvenn web tool (36).

### Minimum free energy (MFE) prediction of miR-mRNA interaction

To assess the thermodynamic stability of the interaction between predicted miRNAs and ID1 3’-UTR, minimum free energy (MFE) of their interaction was calculated, using RNAhybrid web-tool (https://bibiserv.cebitec.uni-bielefeld.de/rnahybrid/) (37). The 3’-UTR sequence of ID1 gene was retrieved from the UTRdb database (http://utrdb.cloud.ba.infn.it/utrdb/) (38) whereas all predicted miRNA sequences were acquired from miRBase (https://www.mirbase.org/) (39). The FASTA sequence of ID1 3’-UTR and individual miRNA sequences was provided as input for MFE analysis. All the analysis were performed using the default parameters of the webtool to achieve the represented MFE values.

### Evaluation of miR-615-5p expression and correlation analysis between miR-615-5p and ID1 in PDAC samples

To analyse the expression profile of miR-615-5p in the clinical samples of Pancreatic Ductal Adenocarcinoma tissues (PDAC), small RNA sequencing data were obtained from Gene Expression Omnibus (GEO) (https://www.ncbi.nlm.nih.gov/gds). The complete set of data were acquired using GSE41369 series. GSE41369 included miRNA expression profile of samples from 9 PDAC patients and 9 normal tissues. The comparative analysis between these two tissue types was performed using GEO2R (http://www.ncbi.nlm.nih.gov/geo/info/geo2r.html), a webtool, integrated with GEO database that allows the comparison of gene expression between two or more experimental groups within a same dataset. Using GEO2R, the TPM normalized values of miR-615-5p from PDAC patients and normal tissues were obtained and bar graph was plotted using GraphPad prism software for statistical analysis. The statistical significance was calculated by Student’s t test (unpaired). To analyse the correlation between miR-615-5p and ID1 expression in PDAC samples, the mRNA expression profile data was obtained from GSE41368 data series. Both these two mentioned sequence data series (GSE41369, GSE41368) was under the same bio-project PRJNA176698 (40). GSE41368 consists of the microarray data depicting the gene expression profiles of 6 PDAC tissues and 6 normal tissues. These samples were also used to analyse the miRNA expression profile in GSE41369. Using GEO2R, the RMA normalized values of ID1 expression from PDAC patients and normal tissues was obtained. Additionally, the common samples from the small RNA-seq dataset (GSE41369) and microarray dataset (GSE41368) were identified using unique GSM identifiers. GSM identifiers are unique for each sample. Spearman correlation analysis (Spearman’s r < 0, p <0.05) was further performed to identify the correlation between miR-615-p and ID1expression in PDAC clinical samples.

### Cell culture and maintenance of cell lines

The human PDAC cell line PANC1 (RRID: CVCL 0480), and human embryonic kidney cell line HEK-293T (RRID: CVCL_0063) were purchased from National repository for cell lines, India (NCCS, Pune). Human Cervical cancer cell line HeLa (RRID: CVCL_0030) was procured from American Type Culture Collection (ATCC, USA). The cell lines mentioned above were maintained and propagated by culturing in Dulbecco’s modified Eagle’s medium (Gibco, Cat. no. 12800017) supplemented with 10% foetal bovine serum, 100 U/ml penicillin and streptomycin (Invitrogen, Cat. no.15140122) at 37 °C in a humidified chamber supplemented with 5% CO_2_. Cell lines were source authenticated using STR profiling and were regularly monitored for contaminations.

### Sequence verification of miR-615-5p

miR-615-5p was amplified from the total RNA of miR overexpressed samples as described previously. The sequence of the forward primer used for PCR amplification of miR-615-5p is 5’ GGGGGTCCCCGGTGCTCGG 3’ and the reverse primer sequence is 5’ GCGAGCACAGAATTAATACGAC 3’. PCR amplified DNA fragments of 65bp were electrophoretically separated on 2.5 % agarose gels, following which the band corresponding to the amplified DNA fragments was extracted and purified following usual procedures. This clear supernatant contains the desired DNA fragment, which was further precipitated by standard protocols, washed with 70% ethanol, air dried, and re-suspended in 15μl TE buffer. The concentration of the eluted DNA was measured and 5ul of the eluted product was used for subsequent cloning into “pJET1.2/blunt Cloning Vector” following manufacturers protocol (Thermo fisher Scientific). The bacterial colonies were then propagated, and plasmids were isolated using QIAprep Spin Miniprep Kit (Qiagen). The isolated plasmids were sequence validated by Sanger sequencing method using vector specific forward primer, supplied with the cloning kit (Thermo fisher Scientific).

### Vectors and transfection

miR-615-5p mimics (C-301123-01-0005), miR-615-5p AntimiR (IH-301123-02-0005), and the corresponding negative control (CN-001000-01-05) were procured from Horizon (USA). The ID1 overexpression construct pcDNA3-hId1 was generously provided by Robert Benezra (Addgene plasmid #16061; RRID: Addgene_16061), pRSpuro shID1#6 knockdown vector was a gift from Joan Massagué (Addgene plasmid #19165; RRID: Addgene_19165), while the FLAG-Ago2 was a gift from Edward Chan (Addgene plasmid #21538, Addgene_21538). For miRNA transfection experiments, HeLa and PANC1 cells were seeded in 6-well plates at densities of 0.8 × 10⁵ and 1.2 × 10⁵ cells per well, respectively, to achieve approximately 50-60% confluency at the time of transfection. Cells were transfected with miR-615-5p mimic at final concentrations of 15 nM (HeLa) and 20 nM (PANC1) using Lipofectamine RNAiMAX (Invitrogen; Cat. no. 13778150). For inhibition studies, both cell lines were transfected with 25 nM miR-615-5p specific AntimiR using the same reagent, following the manufacturer’s recommended protocol. For rescue experiments, cells were first transfected with miRNA mimics and subsequently subjected to plasmid transfection. Briefly, HeLa and PANC1 cells were seeded as described above and, miR-615-5p mimic-transfected cells were rescued by co-transfecting with either 500 ng or 750 ng of pcDNA3-hId1 plasmid, using Lipofectamine LTX (Invitrogen; Cat. no. 15338100). Similarly, for experiments with AntimiR treatment, HeLa and PANC1 cells were transfected first with AntimiR, followed by co-transfection with 1µg of pRSpuro shID1#6 vector using Lipofectamine LTX according to the manufacturer’s instructions. Total RNA and protein lysates were harvested 48 hr post-transfection, unless otherwise specified, and used for downstream qRT–PCR and western blot analyses as required by the experimental design.

### RNA isolation and cDNA synthesis

Total RNA was extracted using Trizol reagent (Ambion, Cat. no. 10296010) following manufacturer’s protocol. To prepare cDNA for mRNA expression analysis, 1μg of total RNA was converted to cDNA using Verso cDNA synthesis kit (Thermo Fisher Scientific, Cat. no. AB-1453/A). For the analysis of miRNA expression,1μg of total RNA was poly-adenylated by incubating the total RNA with poly-A polymerase (Invitrogen, Cat. no. AM2030) and ATP (New England Biolabs, Cat no. P0756S) at 37°C for 1hr. Poly-adenylated RNA was then further converted to cDNA using poly-T specific reverse primer. For the synthesis of cDNA, Super Reverse Transcriptase MuLV Kit (Bio Bharati Life Science, Cat no. BB-E0043) was used. Real-time PCR was further performed using PowerUp SYBR master mix (Applied Biosystems, Cat. no. A25741) on 7500 Fast real-time PCR system (Applied Biosystems, RRID:SCR_018051). All mRNA and miRNA quantification data were normalized to GAPDH and U6 respectively as the case may be. All the primers were procured from Integrated DNA Technologies and listed in the Supplementary Table S7.

### Western immunoblotting

Cells were washed with ice cold PBS and then trypsinised using 0.05% Trypsin-EDTA solution (Invitrogen, Cat. no. 15400054). Harvested cells were then lysed in RIPA buffer (Pierce, Cat no. 89900) supplemented with protease inhibitor cocktail (Sigma Aldrich, Cat. no. P8340) on ice for 1-1.5 hrs. Lysates were then separated from cellular impurities by centrifugation at 14000rpm for 20 minutes at 40C, and protein concentration of the cell lysates were then quantified by BCA method. 40µg-50µg of the total protein was separated on 10% or 15% PAGE and transferred onto PVDF membrane using wet electrophoretic transfer. The membranes were subsequently blocked with 5% BSA and incubated overnight at 4°C with primary antibodies (Anti-ID1: Abcam, Cat. no. ab283650; Anti-LC3B: Abclonal, Cat. no. A19665, RRID: AB_2862723; Anti-Flag: Sigma, Cat. no. F1804, RRID: AB_262044, Anti-GAPDH: Cell Signalling Technology, Cat. no. 2118S, RRID: AB_561053, Anti-β Actin: Cell Signalling Technology, Cat. No. 4967S, RRID: AB_330288). The following day, the membranes were washed 4 times and probed with specific secondary antibodies (Anti-rabbit IgG: Cell Signalling Technology, Cat. no. 7074S, RRID: AB_2099233; Anti-mouse IgG, Cat. no. 7076S, RRID: AB_330924). Membranes were incubated with secondary antibodies for 1hr at room temperature and then visualized with chemiluminescent reagents. Images were acquired with iBright CL-1500 Chemi-luminescent imaging system (Thermo Fisher Scientific, RRID: SCR_026565). The densitometric analyses were carried out using ImageJ (RRID:SCR_003070) and normalized against corresponding loading control. Finally, the intensity of each experimental condition was normalized to the control, and the statistical significance was calculated using GraphPad Prism software (RRID:SCR_002798).

### Immunofluorescence staining and microscopy

HeLa cells were seeded and transfected as described previously. 24hrs post transfection, 0.5 x 10^5^ transfected cells were re-seeded on a glass coverslip in a 12 well plate for 24 hrs, following which the cells were fixed with 1ml of 3.7% pre-warmed paraformaldehyde for 25 mins at room temperature. The fixed cells were then washed for 5 mins with 1X PBS for 3 times and permeabilized with 1ml of 0.1% Triton-X dissolved in 1X PBS for 5 mins. Permeabilized cells were then washed for 3 times and blocked with 1ml of 3% BSA dissolved in 1X PBS for 1 hr at room temperature. Blocked cells were incubated with ID1 specific primary antibody at 1:100 dilution at 4°C in a humidified chamber overnight, following which cells were washed 3 times with 1X PBS and incubated with anti-rabbit Alexa-fluor 488 tagged secondary antibody (Invitrogen, Cat. no. A-11008, RRID: AB_143165) at 1:500 dilution for 1 hr at room temperature. Following the incubations, cells were washed 5 times with 1X PBS and stained with 1ml of the DAPI solution for 5 mins at a concentration of 1: 10000 (Stock conc.:1mg/ml). Cells were then washed 3 times and mounted on a grease free slide using ProLong Antifade Diamond mounting media (Invitrogen, Cat. no. P36965). The mounted coverslip was then photographed using Leica Stellaris DM8 at 63x magnification. To calculate the Mean Fluorescence Intensity (MFI), total fluorescence intensity of each microscopic field was evaluated using ImageJ software (RRID:SCR_003070) and statistical significance of average mean intensity of different conditions were assessed using GraphPad Prism (RRID:SCR_002798). The final represented graph was generated using Biorender (RRID: SCR_018361).

### Dual luciferase assay

To experimentally confirm the interaction between miR-615-5p and ID1 3’-UTR, 68bp fragment of the 3’-UTR encompassing the putative seed sequence and a mutated sequence containing the mutation (GGACCCC to GGAGGGG) in seed region was synthesized and cloned in the PGL3-Basic vector using Xba1 restriction enzyme (New England Biolabs, Cat. No. R0145T), downstream of the firefly luciferase gene to generate the wild-type (WT) and mutated (Mut) construct respectively following the published ((41). Luciferase constructs were sequence verified using EBV reverse primer to ensure the successful insertion of the sequences. The sequence of the UTR region is provided in the Supplementary Table S7. pRL-CMV, ensures a continuous expression of Renilla luciferase and was used for internal normalization of the firefly luciferase activity.

3×10^4^ HEK 293T cells were seeded in a 12 well plate a day before transfection. 125ng wild type or mutant ID1 3’-UTR, both cloned in PGL3-basic vector and 10ng pRL-CMV were co-transfected with either 15nM of miR-615-5p mimic or 25nm of AntimiR or equal concentration of scrambled according to the requirement of the experiments using Lipofectamine 3000. 48 hours post transfection, cells were washed with phosphate-buffered saline (PBS) and subsequently lysed with 1X passive lysis buffer supplied with the Dual-Luciferase Assay kit (Promega, Madison, WI). Total protein concentration in the lysates was measured by BCA Protein Assay (Thermo Fisher Scientific) and equal amount of protein was diluted for each experimental condition was used to measure the luciferase activity following the manufacturer’s protocol. Luminescence was measured by Varioskan Flash multimode reader (Thermo Fisher Scientific). The firefly luciferase activity was normalized against Renilla activity and luciferase activity of different experimental condition was compared to the control condition. Statistical significance of relative luciferase activity was calculated using GraphPad Prism software (RRID:SCR_002798) and the representative graph was generated using Biorender (RRID: SCR_018361).

### AGO2 RNA immunoprecipitation (RIP)

5×10^5^ HeLa cells were seeded onto a 10 cm dish in 10 ml DMEM media overnight, following which 3μg of Flag-Ago2 construct was transfected using Lipofectamine-LTX reagent in Opti-MEM (Gibco, Cat. no. 31985070), for 6-hour, prior to addition of fresh media. The following day, cells were transfected with either 15nM miR-615-5p mimic or 20nM AntimiR or equivalent scrambled controls in Opti-MEM depending on experimental requirement, using Lipofectamine RNAiMax (Invitrogen) for 6 hours, followed by addition of fresh media overnight. Thereafter, 30μl Protein G dyna beads (Invitrogen, Cat. no. 10004D) were prepared for each experimental condition by washing four times with 750μl RIP lysis buffer (150mM Tris-HCl at pH 7.4, 150mM NaCl, 0.5M EDTA, 0.5% NP-40, 5% glycerol and 1x protease inhibitor). The beads were then incubated with 5μg monoclonal anti-Flag (Sigma, Cat. no. F1804, RRID: AB262044) and mouse IgG isotype control (Santa Cruz Biotechnology, Cat. no. sc-2025, RRID: AB_737182) in 500μl RIP lysis buffer overnight for efficient binding. Antibody coated beads were washed 3 to 4 times in 500μl RIP lysis buffer to remove the unbound antibodies and kept at 4°C. 48 hours after the transfection, the cells were first rinsed twice with chilled PBS, scraped and collected by centrifuged at 4500 rpm for 5 mins in 4°C. The cells were then lysed on ice for 45 min using a freshly prepared lysis buffer supplemented with 2-unit Ribolock RNase inhibitors/mL lysis buffer (Thermo Fisher Scientific, Cat. no. EO0382). The cell lysates were then centrifuged at 14,000g at 4°C for 20 min, and the supernatants were collected for the downstream application. The protein concentration in the lysate was measured using BCA reagent (Thermo Fisher Scientific). 400μg of total isolated protein was used for the Co-IP with either Flag or IgG coated Protein G beads for 4 hours at 4°C. 7.5% of the total volume was kept aside separately as input for RNA and protein expression analysis. Following immunoprecipitation, the beads were washed with 500μl RIP lysis buffer 5-7 times. and 20% of the bead was separated from the total volume for western immunoblotting whereas, remaining volume was resuspended in 500μl of RIP lysis buffer. The beads were then incubated with 2.5 U DNase-I (Thermo Fisher Scientific, Cat. no. EN0521) for 10 minutes at 37°C. It was subsequently washed once with 500 μL RIP buffer and incubated with 2.5 U Proteinase K (New England Biolabs, Cat. no. P8107S) for 15 mins at 45°C. Finally, the beads were washed once with RIP lysis buffer and subjected to RNA isolation using TRIzol LS (Invitrogen) following manufacturers protocol. 300ng of eluted RNA was used for ID1 mRNA enrichment analysis in eluted RNA using Verso cDNA synthesis kit (Invitrogen). The miR-615-5p enrichment was analysed by following previously mentioned protocol.

### Biotinylated RNA immunoprecipitation (RIP)

HeLa cells were seeded as mentioned previously. Cells were transfected with 15nM of biotinylated miR-615-5p mimic or biotinylated scrambled for 6 hours using Lipofectamine RNAiMax (Invitrogen) following manufacture’s protocol. 48 hours post-transfection, the cells were washed and lysed following the previously described method. The protein concentration in the lysate was measured using BCA reagent (Thermo Fisher Scientific). 200μl of total isolated protein was used for each immunoprecipitation. For each experimental condition, 30μL of magnetic Streptavidin beads (Thermo Fisher Scientific, Cat. no. 88816) were activated by washing twice with 1ml of 1X washing and binding buffer (5mM Tris-HCl at pH 7.4, 0.5mM EDTA, 1M NaCl), followed by washing with 1 ml of solution B (0.1M NaCl). The beads were further washed and resuspended in 500μL RIP lysis buffer. The activated beads were then blocked with 5μL of BSA (10mg/ml) and 5μL of yeast tRNA (10 mg/ml) (Invitrogen, Cat. no. AM7119) for 30 minutes at 4°C. Bead were then washed and pull-down assay was performed as described previously. The miR-615-5p enrichment was analysed by following previously mentioned protocol.

### Ex-vivo RNA immunoprecipitation (RIP)

5×10^5^ HeLa cells were seeded into a 10 cm dish and was transfected with 3μg of Flag-Ago2 construct using Lipofectamine-LTX reagent in Opti-MEM (Gibco, Cat. no. 31985070) for 6 hrs. 48 hours post transfection cells were processed as described previously. 500ug of total lysate is then incubated with each experimental concentration of AntimiR (1nM, 10nM, 100nM, 200nM) and 200nM of the scrambled control for 30 mins in room temperature and subjected to co-IP. Protein G dyna beads (Invitrogen, Cat. no. 10004D) were washed and incubated with 5μg monoclonal anti-Flag (Sigma, Cat. no. F1804, RRID: AB262044) and mouse IgG isotype for overnight at 4°C. The pull-down was performed using the previously described process. Following RNA isolation 300ng of eluted RNA were used for ID1 mRNA and miR-615-5p enrichment analysis following above mentioned protocol

### In-vitro Migration assay

For trans-well migration assay, 0.8 × 10^5^ HeLa and 1.2 × 10^5^ PANC1 cells were grown on 6-well culture plates to attain 50–60% confluency. For ID1 complementation experiment, 15nM and 20nM of miR 615-5p mimic were transfected in HeLa and PANC1 cells respectively using Lipofectamine RNAiMax. Over-expression of ID1 is achieved by transfecting 500 ng and 750ng of pcDNA3 hId1 in HeLa and PANC1 cells respectively using Lipofectamine LTX. For ID1 knockdown experiment, 24hrs post-seeding 25nM of AntimiR in both the cell lines were transfected using Lipofectamine RNAiMax, followed by co-transfection with 1µg of pRSpuro shID1#6 plasmid using Lipofectamine LTX. All transfections were performed in Opti-MEM for 6 hours. 24 hrs post transfection, cells were washed with 2ml of pre-warmed (37°C) PBS and then trypsinised with 500 μl of 0.05% Trypsin-EDTA solution for 2-3 minutes following which cells were suspended and washed 2 times with 5ml serum free media (SFM). Cells were then counted and 1 × 10^5^ HeLa or 0.5× 10^5^ PANC1 were placed in the upper chamber of the cell culture inserts with 300μl serum free medium. The cells were allowed to migrate for 24-30 hrs in response to complete medium (1ml) placed in the bottom chamber of the 12-well trans-well plates. Cells that migrated to the lower surface of the membrane were fixed with 3.7% formaldehyde, stained with 0.5% crystal violet. The non-migrated cells were cleaned carefully and at least 15 independent microscopic fields were counted under microscope. Average no. of migrated cells was counted ImageJ software (RRID:SCR_003070) and comparative bar graph was generated, and statistical significance was calculated using GraphPad Prism (RRID:SCR_002798). The represented graph was generated using Biorender (RRID: SCR_018361).

### Autophagy Flux assay

To assess the autophagic flux, 0.8 × 10^5^ HeLa and 1.2 × 10^5^ PANC1 cells were grown on 6 well culture plates. Following attainment of 50-60% confluency, mimic or AntimiR (at concentrations mentioned previously) were transfected for 6hrs in Opti-MEM using Lipofectamine RNAiMAX following the manufacturer’s protocol, followed by addition of fresh growth medium post 6 hrs of transfection. 46 hrs post transfection, cells were treated with 50μM CQ (for 2 hrs) to attain autophagic block. Cells were then washed with ice cold PBS and scraped on ice, lysed using RIPA lysis buffer and subjected to western immunoblotting to assess the expression of LC3B-II. Relative LC3B-II expression was analysed using ImageJ software (RRID:SCR_003070) and normalized against corresponding GAPDH with respect to Control. To assess the autophagic flux in different experimental conditions, relative change of LC3B-II level upon CQ treatment was compared with CQ untreated condition (LC3B-II_CQ Treated_ –LC3B-II_CQ Untreated_) for each experimental condition. The difference represents autophagy flux for each experimental condition. The flux in mimic and AntimiR treated condition was then normalized against the flux in Scr control to assess the changes in different experimental condition as compared to the Scr control. Statistical significance of the average flux between different experimental conditions was evaluated using GraphPad Prism (RRID:SCR_002798) and represented graph was generated using Biorender (RRID: SCR_018361).

### Statistical analysis

Student’s t-test (paired and unpaired) was used to determine the p-values using GraphPad Prism software (Version 5.01) (RRID:SCR_002798). Graphs were prepared using Biorender (RRID: SCR_018361). In each case, the value of (ns) p > 0.05, (*) p ≤ 0.05, (**) p ≤ 0.01, and (***) p ≤ 0.001 were considered statistically significant.

## Results

### ID1 is overexpressed in several cancers and is associated with poor prognosis in PDAC

To assess the oncogenic relevance of Inhibitor of Differentiation 1 (ID1) across human malignancies, we performed a pan-cancer *in silico* analysis using the GEPIA2.0 platform, integrating transcriptomic data from TCGA and GTEx cohorts. ID1 expression was significantly elevated in multiple cancer types, including Diffuse Large B-cell Lymphoma (DLBC), Glioblastoma Multiforme (GBM), Lower Grade Glioma (LGG), Pancreatic Adenocarcinoma (PAAD), Testicular Germ Cell Tumours (TGCT), and Thymoma (THYM), indicating a broad oncogenic association of ID1 across histologically diverse tumours (**Figure 1a**). Survival analysis using the Kaplan-Meier Plotter revealed that high ID1 expression was significantly associated with reduced overall survival (OS) and disease-free survival (DFS) in PAAD patients (log-rank p < 0.05; **Figure 1b-c**), suggesting a prognostic role for ID1 in pancreatic cancer. Hazard ratio (HR) analysis revealed a 32% (in case of OS) and 38% (in case of DFS) increased risk of mortality and disease recurrence, in PAAD patients with high ID1 expression. Given that pancreatic ductal adenocarcinoma (PDAC) constitutes over 90% of PAAD cases and represents the most aggressive pancreatic cancer subtype, we further examined ID1 expression specifically in PDAC. Analysis of TCGA-PDAC samples demonstrated a greater than three-fold upregulation of ID1 in tumour tissues compared to normal pancreas (**Figure 1d**). Consistently, elevated ID1 expression in PDAC patients was also associated with significantly reduced overall survival (log-rank p < 0.05), with hazard ratio analysis indicating a 58% increased risk of mortality in the high-ID1 expression group **(Figure 1e)**. Collectively, these data along with previous reports on the oncogenicity of ID1 establish ID1 as a robust prognostic indicator and supports its potential role as a contributor of tumour progression, particularly in PDAC.

**Figure 1:**
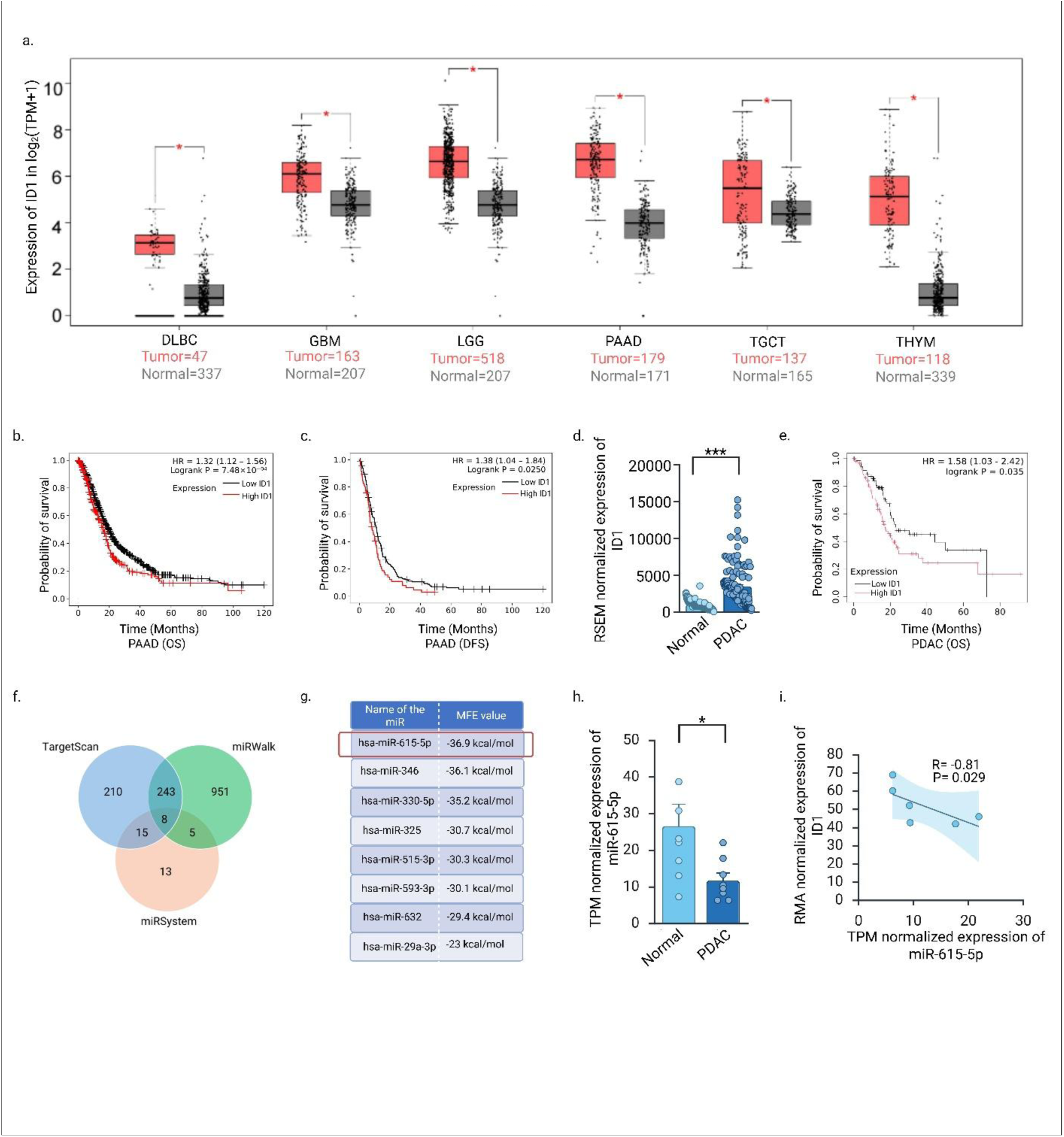
ID1 is overexpressed in several cancers and is associated with poor prognosis in PDAC. In silico analysis using GEPIA 2.0 shows that ID1 expression is significantly upregulated in Diffused Large B-cell Lymphoma (**DLBC**), Glioblastoma Multiforme (**GBM**), Lower Grade Glioma (**LGG**), Pancreatic Adenocarcinoma (**PAAD**), Testicular Germ Cell Tumors (**TGCT**) and Thymoma (**THYM**). The screen parameters were as follows: p<0.05, cutoff of |Log_2_FC|=1. The statistical significance was analyzed by using one way ANOVA. b-c) The overall survival (OS), disease free survival (DFS) probability was analyzed using Kaplan-Meier Plotter. A significantly reduced probability of overall survival and disease-free survival (DFS) was observed in High ID1 expressing PAAD groups (p<0.05). The Hazards Ratio (HR) suggested 32% lower risk of overall survival and 38% lower risk of disease free among high ID1 expression groups. d) Expression of ID1 in PDAC was acquired from TCGA and compared with its expression in normal pancreas retrieved from GTEX using UCSC XENA. UCSC XENA uniformly processes the datasets and represents as RSEM normalized values, which allows cross-cohort comparison of gene expressions. The result shows a significant up-regulation of ID1 expression in PDAC samples as compared to normal. ID1 expression was compared between 150 PDAC samples and 167 normal tissues and ∼ 3-fold increased expression was observed in PDAC tissues (p< 0.0001). e) Overall survival of high ID1 expressing groups of PDAC patients was analyzed by Kaplan-Meier Plotter. Elevated expression of ID1 is associated with significantly reduced overall survival among PDAC patients (p < 0.05), with hazard ratio analysis indicating a 58% increased risk of mortality in the high-ID1 expression PDAC groups. f) ID1 targeting microRNAs were identified using TargetScan, miRWalk, miRSystem, which predicted 476, 1207, 41 ID1 targeting miRs respectively. To minimize false positivity, a comparative analysis was carried out manually. Comparative analysis of the predicted microRNAs from 3 different sets of prediction tools identified 8 common microRNAs, namely, hsa-miR-615-5p, hsa-miR 346, hsa-miR-330-5p, hsa-miR-325, hsa-miR-515-3p, hsa-miR-593-3p, hsa-miR 632, has-miR-29a-3p. g) Minimum free energy (MFE) of the interaction between ID1 3’UTR and predicted microRNAs was calculated using RNAhybrid web-tool which identified miR-615-5p/ID1 interaction as most thermodynamically stable. The table represents the MFE values (kcal/mol) for each predicted microRNA and ID1 interaction. Considering the MFE values, hsa-miR-615-5p/ ID1 binding exhibits lowest MEF (–36.9 kcal/mol) among all. h) The expression of miR-615-5p is significantly downregulated in PDAC clinical samples compared to the normal adjacent tissues. Small RNA sequencing data of the PDAC clinical tissue samples were analyzed from GSE41369 series using GEO2R. TPM normalized expression values of miR-615-5p from PDAC tumor and normal tissues were represented as bar graph. TPM or transcripts per million is a commonly used method to normalize RNA-seq data and allows the gene expression levels comparable both within and between the samples. During RNA sequencing, different genes produce different numbers of reads depending on their length and the sequencing depth of each sample. TPM normalization corrects these biases and represents the expression of a gene as the number of transcripts present in 1 million total reads. The data represented as average ± SE and significance was calculated using unpaired Students t-test (*: p <0.05). i) ID1 expression shows an inverse correlation with miR-615-5p expression in PDAC clinical tissues. mRNA expression data was obtained from GSE413368 data series. The microarray data was analysed using GEO2R and RMA normalized expression values of ID1 transcript were obtained and used for correlation analysis. RMA or Robust Multi-array Average is a widely used normalization method for the processing of microarray gene expression data from Affymetrix platforms. It corrects technical variations to make expression levels comparable across samples by background correction, quantile normalization, and summarization. The output values are log2 transformed, making data easier to interpret and compare across the samples. The TPM normalized expression values of miR-615-5p and RMA normalized expression values of ID1 transcript of the identical samples were tested for Spearman’s correlation analysis. The result depicts a significantly negative correlation between miR-615-5p and ID1 expression in PDAC tumor tissues (Spearmen’s r < 0; p < 0.05).

### miR-615-5p negatively correlates with ID1 expression in PDAC clinical samples

To identify microRNAs (miRNAs) that may regulate ID1 expression, an integrative *in silico* analysis was performed using three independent target prediction platforms: TargetScan, miRWalk, and miRSystem. To enhance prediction stringency, only miRNAs consistently identified across all three algorithms were shortlisted. This approach yielded eight candidate miRNAs predicted to target the 3′-untranslated region (3′-UTR) of ID1 mRNA (**Figure 1f**). The predicted miRNA-ID1 interactions were subsequently evaluated based on their thermodynamic stability using RNAhybrid. Minimum free energy (MFE) calculations revealed differential binding affinities among the candidate miRNAs, with more negative MFE values indicating energetically favourable interactions. Among the eight candidates, miR-615-5p exhibited the lowest MFE (–36.9 kcal/mol), suggesting the strongest predicted interaction with the ID1 3′-UTR **(Figure 1g)**. Accordingly, miR-615-5p was prioritized for subsequent analyses.

To assess the clinical relevance of miR-615-5p in pancreatic ductal adenocarcinoma (PDAC), small RNA sequencing data from PDAC tissues (GSE41369) were analysed using GEO2R. miR-615-5p expression was significantly reduced in PDAC samples compared with normal pancreatic tissues (p < 0.05) **(Figure 1h)**. In parallel, analysis of ID1 expression using matched microarray data from the same cohort (GSE41368) revealed a significant inverse correlation between miR-615-5p and ID1 expression levels, as determined by Spearman correlation analysis (Spearman’s r < 0, p < 0.05) **(Figure 1i)**. Collectively, these results not only confirm an inverse association between the tumour suppressor miR-615-5p and the oncogene, ID1 in PDAC, at the same time strongly suggest a plausible negative regulation of the oncogene by the miR-615-5p.

### ID1 gene expression is negatively regulated by miR-615-5p

*In silico* data indicated an inverse relationship between miR-615-5p and ID1 expression suggesting a plausible regulation of ID1 by miR-615-5p. Based on *in silico* target prediction and inverse expression patterns observed in PDAC clinical datasets, the regulatory relationship between miR-615-5p and ID1 was experimentally investigated using gain and loss-of-function approaches. Ectopic overexpression of miR-615-5p using miR mimic significantly reduced ID1 protein levels in PANC1 and HeLa cells, as determined by western blotting **(Figure 2a-b)** and immunofluorescence analyses **(Figure 2g)**. Densitometric data revealed a significant reduction in ID1 expression (PANC1= ∼ 50%, p <0.05; HeLa= ∼ 60%, p <0.0001) relative to control conditions, as shown in **Supplementary fig. S2a-b**. Quantified data from the immunofluorescence analysis **(Figure 2g)** shows a significant decrease in the expression of ID1 ( ∼ 70%, p= <0.0001) as indicated by the cumulative Mean Fluorescence Intensity (MFI) graph in the cells overexpressing miR-615-5p. A comparable decrease in ID1 expression was also observed in the glioblastoma cell line CCF52 following miR-615-5p overexpression **(Figure 2c)**, with reduction in ID1 levels by ∼70% (p< 0.02) (**Supplementary fig. S2c)**, indicating that this regulatory effect is conserved across distinct cellular contexts. Conversely, functional inhibition of endogenous miR-615-5p using a sequence-specific AntimiR led to a significant upregulation of ID1 expression in PANC1, HeLa and MIA PaCa-2 cells (**Figure 2d-f**). The expression level of miR-615-5p upon over-expression with miR-mimic, and AntimiR transfection in the mentioned cell lines were shown in **Supplementary fig. S1a-c** and **Supplementary fig. S1d-f** respectively. ID1 protein levels increased by 1.8 to 2-folds (PANC1= ∼ 1.8-Fold, p= <0.0001, HeLa= ∼ 2.6-Fold, p <0.0001, MIA PaCa-2= ∼ 1.8-Fold, p= 0.009) upon miR-615-5p inhibition compared with respective controls, as indicated by densitometric data **(Figure 2d-f, Supplementary fig. S2d-f)** as well as MFI data (∼ 2-Fold, p <0.0001) **(Figure 2h)**, supporting a suppressive role for miR-615-5p in regulating ID1 expression. To further establish the specificity of this inverse association between miR-615-5p and ID1 expression, rescue experiments were performed in which miR-615-5p mimics were reintroduced in both HeLa and PANC1 cells in the presence of the AntimiR. Re-expression of miR-615-5p effectively reversed the AntimiR-induced upregulation of ID1 dose-dependently in both PANC1 **(Figure 2i)** and HeLa cells **(Figure 2j)**. ID1 expression decreased progressively (PANC1 decrease= >60% at 20nM, p <0.05, >65% at 25nM, p <0.005; HeLa decrease= >60% at 15nM, p =0.005; >70% at 20nM, p =0.0001, >75% at 25nM, p <0.0001) with increasing concentrations of miR-615-5p mimic, which is evident from the densitometric analysis **(Supplementary fig. S2g-h)**, confirming that ID1 regulation is specifically mediated by miR-615-5p. Together, these findings support miR-615-5p negatively regulates ID1 expression which aligns with our previous *in silico* data.

**Figure 2:**
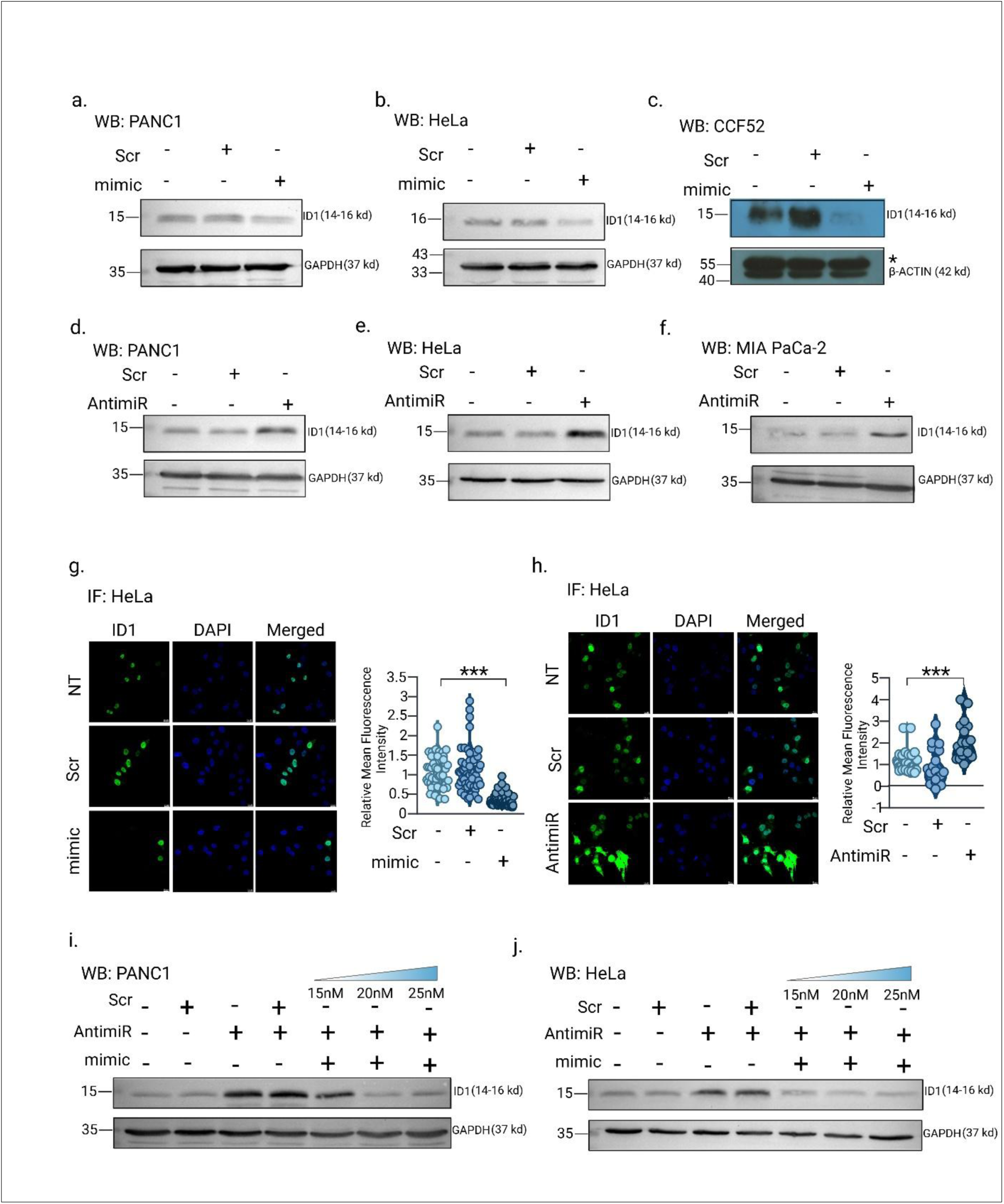
miR-615-5p negatively regulates ID1 expression. **a-c**) Ectopic overexpression of miR-615-5p by transient transfection of miR-615-5p specific synthetic oligos (mimic) led to downregulation of ID1 in PANC1, HeLa, and CCF52 cell line. a and b are the representative of at least 3 independent experiments, whereas, c is the representative of 2 similar repeats. The asterisk (*) indicates a non-specific band detected by the β-actin antibody and does not correspond to the target protein. 15nM, 20nM, 32.5nM of the mimics were used for overexpressing miR-615-5p in HeLa, PANC1 and CCF52 respectively. d-f) Inhibiting miR-615-5p function by transfecting sequence specific miR-615-5p AntimiR upregulates ID1 expression in PANC1, HeLa and MIA PaCa-2 cell lines. The expression of ID1 was evaluated by western immunoblotting. To block miR-615-5p function, 25nM of the AntimiR was transfected in PANC1, HeLa and MIA PaCa-2 cells. The image is a representative of 3 independent experiments. g) Immunofluorescent staining upon ectopic overexpression of miR-615-5p significantly downregulates (∼ 70%) ID1 gene expression. The mean fluorescence intensity was calculated using ImageJ software and normalized against non-transfected control (NT). At least 10 microscopic fields per experimental condition were considered to assess the mean intensity. The violin plot represents mean fluorescence intensity ± SE of 4 independent experiments and significance was calculated using paired Students t-test, ***P < 0.001. h) Immunofluorescent staining upon suppression of miR-615-5p by sequence specific AntimiR led to the ∼ 2-Fold upregulation of ID1 expression in HeLa cells. The mean fluorescence intensity was calculated using ImageJ software and normalized against non-transfected control (NT). At least 10 microscopic fields per experimental condition were considered to assess the mean intensity. The violin plot represents mean fluorescence intensity ± SE of 2 independent experiments and significance was calculated using paired Students t-test, ***P < 0.001. i-j) Overexpression of miR-615-5p competitively reverses AntimiR-mediated ID1 upregulation highlighting the specificity of ID1 regulation by miR-615-5p in both PANC1 and HeLa cells, respectively. Elevated ID1 expression upon inhibiting miR-615-5p was reversed by subsequent reintroduction of miR-615-5p. ID1 expression was evaluated by Western immunoblotting. The data demonstrated that introduction of AntimiR (25nM) caused an increased ID1 expression which was reversed upon dose dependent overexpression of miR-615-5p (15nM, 20nM, 25nM) in PANC1 and HeLa cells. The image is a representative of 3 independent experiments.

### miR-615-5p binds to the GGACCCC binding site within the 3′-UTR of ID1

*In silico* analysis using TargetScan predicted a putative miR-615-5p binding site within the 3′-UTR of the ID1 transcript. To experimentally validate this predicted binding, a luciferase reporter construct containing the wild-type ID1 3′-UTR fragment encompassing the putative binding site and a reporter construct containing mutation in the miR-615-5p binding site were generated **(Figure 3a)**. Co-transfection of the wild-type ID1 3′-UTR reporter with miR-615-5p mimic in HEK293T cells resulted in a significant reduction in luciferase activity compared to scrambled control as indicated by a significant decrease in reporter activity by ∼ 60% (n = 7, p < 0.0001), as shown in **Figure 3b**, suggesting that miR-615-5p binding to the 3-UTR of ID1 suppress its transcriptional activity. Unlike the wild-type reporter, co-transfection of the mutant construct with miR-615-5p mimic did not result in any reduction in the luciferase activity **(Figure 3b)**, demonstrating that the binding of predicted seed-binding site to the 3′-UTR of ID1 is necessary for miR-615-5p mediated repression of the ID1. To further validate this interaction, a sequence-specific AntimiR was co-transfected with the wild-type ID1 3′-UTR reporter in HeLa cells. Inhibition of miR-615-5p led to a significant increase in luciferase activity compared with control conditions **(Figure 3c)**, with reporter expression increased by >2.5 fold (n = 6, p = 0.008). In contrast, AntimiR-mediated inhibition of miR-615-5p failed to enhance luciferase activity of the mutant reporter construct, indicating that ID1 carries a putative binding site at its 3′-UTR for miR-615-5p and can be directly regulated by miR-615-5p.

**Figure 3:**
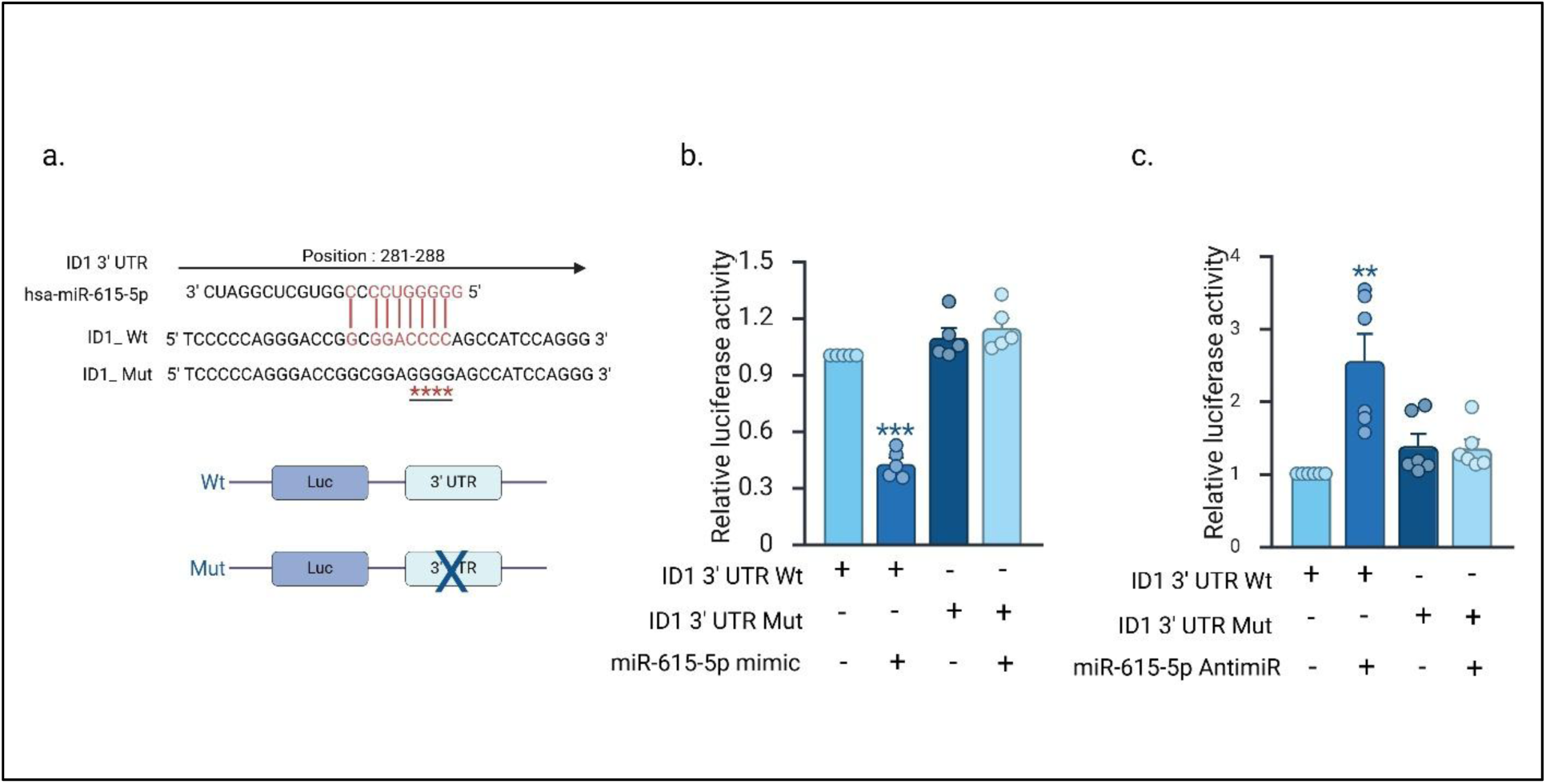
Functional validation of miR-615-5p binding site within the 3’UTR of ID1 gene by Luciferase assay. **a**) A schematic diagram representing the position of miR-615-5p binding site in the 3’UTR region of ID1 gene as predicted by TargetScan. The interacting sequences between miR-615-5p seed region and the wildtype ID1 3’UTR (Wt) are marked in red. The mutation in miR-615-5p seed recognition site in 3’UTR (Mut) is marked in the diagram (*). The 3’UTR sequences were cloned at the 3’ of Firefly luciferase expressing PGL3-basic vector and used for luciferase assay. b) Relative luciferase activity of Wt and mutated construct of ID1 3’UTR (125ng) was assessed upon miR-615-5p overexpression (15nM) in HEK-293T cells. Renilla luciferase expression (10ng) was used in internal normalization. A ratio of firefly and renilla luciferase activity was normalized against scrambled (Scr) control (15nM). The data represented as average ± SE of 7 independent experiments and significance was calculated using paired Students t-test (***p <0.0001). c) Relative luciferase activity of Wt and mutated construct of ID1 3’UTR (125ng) was assessed upon inhibiting miR-615-5p by AntimiR (25nM) in HeLa cells. Renilla luciferase expression (10ng) was used for internal normalization. A ratio of firefly and renilla luciferase activity was normalized against scrambled (Scr) control. The result was represented as average ± SE of 6 independent experiments and significance was calculated using paired Students t-test (**p <0.001).

### miR-615-5p interacts with the 3’-UTR of ID1 mRNA and forms complex with AGO2

To investigate the molecular basis underlying miR-615-5p mediated repression of ID1, RNA immunoprecipitation (RIP) based approaches were undertaken to determine whether miR-615-5p physically interacts with the ID1 transcript and associates within AGO2-containing RISC complex *in vitro*. To directly assess whether miR-615-5p associates with the ID1 transcript, HeLa cells were transfected with biotinylated miR-615-5p mimics, and enrichment of ID1 transcript in the streptavidin pulldown samples was assessed by RT-PCR. Quantitative analysis of the recovered pulled down RNA revealed a significant enrichment of ID1 mRNA in the miR-615-5p pulldown **(Figure 4a)** compared with scrambled conditions (ID1= ∼2.9-fold, p = 0.0001, n=4), indicating a direct association between miR-615-5p and the ID1 transcript. Efficient delivery of the miRNA mimic (**Figure 4b-c**) and basal ID1 expression levels were independently confirmed by qRT–PCR, while densitometric analyses of immunoblot results confirmed ∼ 55% downregulation of ID1 expression (n=3, p= <0.0001) respectively (**Figure 4d, Supplementary fig. S3a**). To assess the ability of miR-615-5p to interact with ID1 and subsequent association with RISC complex, AGO2-RIP assays were performed. Immunoprecipitation of FLAG-AGO2 using anti-FLAG antibody resulted in significant co-enrichment of both ID1 mRNA and miR-615-5p relative to the scrambled controls (ID1 enrichment = > 3.2-fold, n=4, p= 0.0009; miR-615-5p enrichment = > 15-fold, n=3, p= 0.00), consistent with the RIP data with biotinylated mimic **(Figure 4e-f)**. Efficiency of AGO2 pulldown **(Figure 4h)** and downregulation of ID1 protein levels (decrease ∼ 50%, n=3, p= <0.0001) **(Figure 4g, Supplementary fig. S3b)** were confirmed by immunoblot analysis. Of note in this AGO2 RIP data, the significant enrichment of ID1 **(Figure 4e)** in pull-down samples for the mimic transfected conditions versus the control, as opposed to the downregulation of ID1 expression **(Figure 4g)** in response to mimic overexpression, is indicative of the fact that functional downregulation of ID1 expression occurs only upon physical association of the ID1 with the AGO2-RISC complex. The specificity of this association was further assessed by functionally inhibiting miR-615-5p using a sequence-specific AntimiR. AntimiR treatment resulted in a marked reduction in the enrichment of both ID1 mRNA **(Figure 4i)** and miR-615-5p **(Figure 4j)** in AGO2 immunoprecipitated samples (ID1 reduction = > 45%, n=4, p= 0.0015, miR-615-5p reduction= ∼ 47%, n= 4, p <0.0001) **(Figure 4i-j)** accompanied by a significant increase in ID1 expression (fold change = >2-Fold, n= 3, p= 0.0002) **(Figure 4k, Supplementary fig. S3c)**. Efficiency of Ago2 pulldown was checked by western blot as shown in **Figure 4l**. To exclude potential nonspecific effects arising from cellular perturbations associated with AntimiR transfection, an *ex-vivo* competition RIP assay was performed under minimally perturbed conditions. Cell lysates were incubated with increasing concentrations of miR-615-5p specific AntimiR (1, 10, 100, 200 nM) prior to AGO2 immunoprecipitation. The outcome revealed a dose-dependent decrease in ID1 transcript enrichment upon AGO2 pulldown (Fold change= >70%, n=3, p <0.0001) with increasing AntimiR concentrations **(Figure 4m), as well as miR-615-5p enrichment (Figure 4n**; Fold change= ∼ 90%, n=3, p<0.0001). Efficiency of AGO2 pulldown across all conditions was confirmed by immunoblotting (**Figure 4o**). Collectively, these RIP-based analyses demonstrate that miR-615-5p directly associates with the ID1 transcript and forms AGO2 complexes, providing mechanistic insight for miR-615-5p mediated post-transcriptional regulation of ID1.

**Figure 4:**
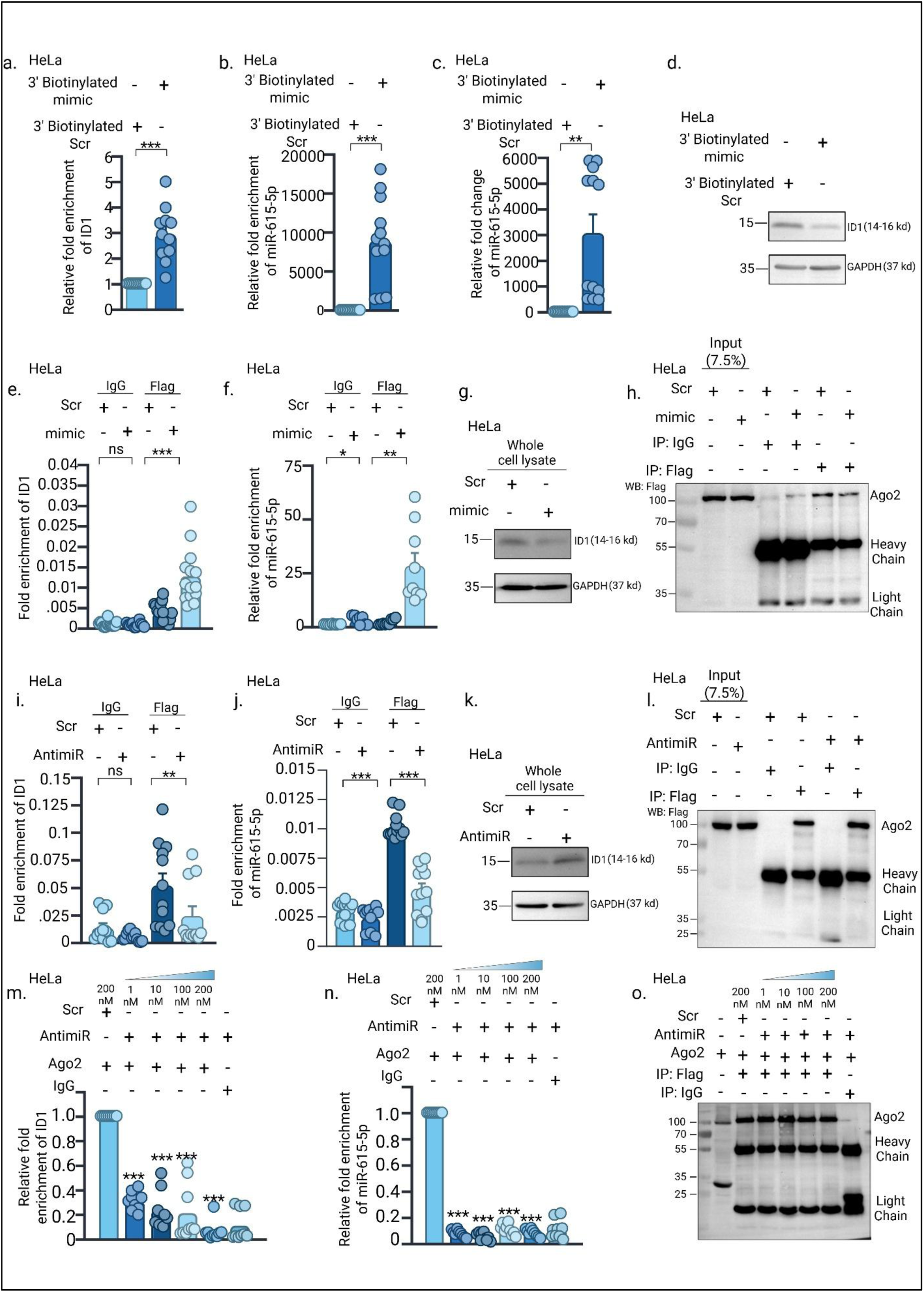
miR-615-5p physically interacts with ID1 3’UTR and forms complex with AGO2. **a**) RNA immunoprecipitation (RIP) assay with 3’ biotinylated miR-615-5p mimic showed a significant enrichment of ID1 transcript upon RIP. ID1 enrichment was assessed by real-time PCR. The expression of ID1 upon 3’ biotinylated miR-615-5p RIP was normalized against 3’ biotinylated scrambled (Scr) control. The data represented as average ± SE of 4 independent experiments and significance was calculated using paired Students t-test (***p <0.0001). b) The bar graph represents significant enrichment of miR-615-5p upon 3’biotinylated miR-615-5p mediated RIP. The expression of miR-615-5p in the pull-down samples was evaluated by real-time PCR. The enrichment of miR-615-5p upon 3’ biotinylated miR-615-5p mediated RIP assay was normalized against the scrambled control (Scr). The data was represented as average ± SE of 4 independent experiments and significance was calculated using paired Students t-test (***p <0.0001). c) The bar graph represented the expression of miR-615-5p upon 3’ biotinylated miR-615-5p transfection. RNA was isolated from the total cell lysate to validate the miR-615-5p overexpression. U6 was used as an internal control, and overexpression was normalized against Scr control. The result was represented as average ± SE of 4 independent experiments and significance was calculated using paired Students t-test (**p <0.001). d) 3’ biotinylated miR-615-5p transfection led to a significant reduction in ID1 expression, as validated by western immunoblotting. The image is the representative of 3 independent experiments. e) AGO2-RIP Assay showed a significant enrichment of ID1 transcript upon miR-615-5p overexpression. ID1 expression in the pulldown sample was evaluated by real-time PCR. To calculate the ID1 enrichment, the Ct value of ID1 fraction (Ct^RIP^) in the pull-down sample was subtracted from the C_t_ value of ID1 mRNA in the 7.5% input sample (Ct^Input^). The linear conversion of ΔCt (Ct^Input^ – Ct^RIP^) was used to assess the enrichment level of ID1 transcript of the pulldown samples. The data was represented as average ± SE of 4 independent experiments and significance was calculated using paired Students t-test (***p <0.0001). f) The result represents the enrichment of miR-615-5p upon Ago2-RIP assay. Enrichment analysis showed a significant association of miR-615-5p with AGO2 upon miR-615-5p overexpression. To calculate miR-615-5p enrichment analysis, the Ct value of miR-615-5p fraction (Ct^RIP^) in the pull-down sample was normalized against Ct value of miR-615-5p fraction from IgG isotype control of Scrambled (Scr) treated condition. The linear conversion of the normalized Ct value was represented as relative fold enrichment of miR-615-5p. The data was represented as average ± SE of 3 independent experiments and significance was calculated using paired Students t-test (*p <0.05, **p <0.001). g) ID1 expression in the whole cell lysate was assessed through western immunoblotting with ID1 specific antibody. GAPDH was used as an internal loading control. The image is the representative of 3 independent experiments. h) The pull-down efficiency of the Flag-AGO2 was assessed by western immunoblotting using primary antibody specific to the Flag-tag. The represented image is a representative of four such images. i) AGO2-RIP Assay showed a significant downregulation in the enrichment of ID1 transcript upon functional inhibition of miR-615-5p by AntimiR. ID1 expression in the pulldown sample was evaluated by real-time PCR. To calculate the ID1 enrichment, the Ct value of ID1 fraction (Ct^RIP^) in the pull-down sample was subtracted from the Ct value of ID1 mRNA in the 7.5% input sample (Ct^Input^). The linear conversion of ΔCt (Ct^Input^ – Ct^RIP^) was used to assess the enrichment level of ID1 transcript of the pulldown samples. The data was represented as average ± SE of four independent experiments and significance was calculated using paired Students t-test (**p <0.001). j) The histogram represents the enrichment level of miR-615-5p upon Ago2-RIP assay in the presence of AntimiR. Enrichment analysis showed a significant downregulation of miR-615-5p with Ago2 upon miR-615-5p inhibition. To calculate miR-615-5p enrichment, the Ct value of miR-615-5p fraction (Ct^RIP^) in the pull-down sample was subtracted from the Ct value of miR-615-5p mRNA in the 7.5% input sample (Ct^Input^). The linear conversion of ΔCt (Ct^Input^ – Ct^RIP^) was used to assess the enrichment level of miR-615-5p in the pulldown samples. The data was represented as average ± SE of four independent experiments and significance was calculated using paired Students t-test (***p <0.0001). k) ID1 expression in the whole cell lysate was assessed through western immunoblotting with ID1 specific antibody. GAPDH was used as an internal loading control. The image is the representative of 3 independent experiments. l) The pull-down efficiency of the Flag-AGO2 was assessed by western immunoblotting using primary antibody specific to the Flag-tag. The represented image is representative of 4 such experiments. m) AntimiR mediated competitive RIP-assay showed a dose dependent decrease in ID1 enrichment with concomitant increase of AntimiR. To calculate the relative fold change of ID1 transcript, the Ct value of ID1 fraction in the pulldown RNA (Ct^RIP^) was subtracted from the Ct value of ID1 in the Input (Ct^Input^). The Δ^Ct^ value was then normalized against Scrambled (Scr) control to achieve ΔΔ^Ct^. Linear conversion of ΔΔ^Ct^ was represented as relative fold enrichment. The data represented as average ± SE of 3 independent experiments and significance was calculated using paired Students t-test (***p <0.0001). n) Introduction of AntimiR prior to competitive RIP-assay shows a significantly decreased expression of miR-615-5p enrichment with concomitant increase of AntimiR concentration. To calculate the relative fold change of ID1 transcript, the Ct value of ID1 fraction in the pulldown RNA (Ct^RIP^) was subtracted from the Ct value of ID1 in the Input (Ct^Input^). The Δ^Ct^ value was then normalized against Scrambled (Scr) control to achieve ΔΔ^Ct^. Linear conversion of ΔΔ^Ct^ was represented as relative fold enrichment. The data represented as average ± SE of 3 independent experiments and significance was calculated using paired Students t-test (***p <0.0001). o) The pull-down efficiency of the Flag-AGO2 was assessed by western immunoblotting using primary antibody specific to the Flag-tag. The represented image is representative of 3 such experiments.

### miR-615-5p suppresses ID1 mediated cellular migration

A negative correlation between expression of miR-615-5p and ID1, or their regulatory interaction doesn’t necessarily imply functional changes in the context of carcinogenesis. To assess whether miR-615-5p mediated repression of ID1 influences cellular migratory behaviour, miR-615-5p was ectopically overexpressed in the PDAC cell line PANC1 **(Figure 5a-b** and **Figure 7d-e),** as well as in HeLa cells **(Figure 5g-h)**. Data clearly depicts that ectopic over-expression of miR-615-5p mimic caused a significant reduction in the number of migrated PANC1 cells (∼ 60%, n= 3, p <0.0001) and HeLa cells ( >70%, n=3, p <0.0001), which was restored upon over-expressing ID1 either in PANC1 **(Figure 5a-b**, and **Figure 7d-e)** or in HeLa cells **(Figure 5g-h)**. Consistent with reduced cell migration upon ectopic over-expression of miR-615-5p in both PANC1 as well as in HeLa cells, miR-615-5p overexpression significantly suppressed ID1 protein levels relative to scrambled controls in both PANC1 **(Figure 5c, Supplementary fig. S4a)** as well as in HeLa **(Figure 5i, Supplementary fig. S4c),** which is restored upon ID1 complementation. These observations indicate that increased miR-615-5p abundance is associated with reduced ID1 expression and suppressed cell migration. To determine the role of ID1 in miR-615-5p induced suppression of migration, rescue experiments were performed by reintroducing ID1 in cells overexpressing miR-615-5p. Re-expression of ID1 significantly reversed the miR-615-5p mediated reduction in migratory capacity (mimic+pcDNA3 vs mimic+ID1= >3.5-fold, n= 3, p <0.0001) **(Figure 5a)**. These results indicate that suppression of cell migration by miR-615-5p is largely dependent on downregulation of ID1. Furthermore, consistent with the tumour suppressive role of miR-615-5p, functional inhibition of miR-615-5p using AntimiR significantly upregulated the *in vitro* migratory ability of both PANC1 (>2-fold, n= 3, p <0.0001) **(Figure 5d-e)** and HeLa cells (∼ 2-fold, n=3, p <0.0001) **(Figure 5m-n)** coupled with increased expression of ID1 (**Figure 5f, Supplementary fig. S4b; Figure 5l, Supplementary fig. S4d; Figure 5o, Supplementary fig. S4e)**. The phenomenon was reversed upon selective knockdown of ID1 in presence of AntimiR (Reversal _PANC1_=>39%, n=3, p= <0.0001; Reversal _HeLa=_ >36%, n=3, p <0.0001) **(Figure 5d-e**, **Figure 5m-n).** Additionally, to validate the specificity of the miR-615-5p/ID1 regulatory interaction in controlling cell migration, competition assays were performed. Inhibition of miR-615-5p using AntimiRs led to a significant increase in migratory capacity of HeLa cells (fold increase = >1.4 fold, n= 2, p= 0.0001) (**Figure 5j-k)**, concomitant with elevated ID1 expression **(Figure 5l, Supplementary fig. S4d)**. Whereas reintroduction of miR-615-5p mimics under these conditions reversed both ID1 expression (**Figure 5l, Supplementary fig. S4d**) and the associated cellular migration (AntimiR+scr vs AntimiR+mimic= >40%, n=2, p <0.0001) (**Figure 5j-k**). These findings demonstrate and confirm that miR-615-5p dependent modulation of cell migration is specifically mediated through its regulation of ID1 expression.

**Figure 5:**
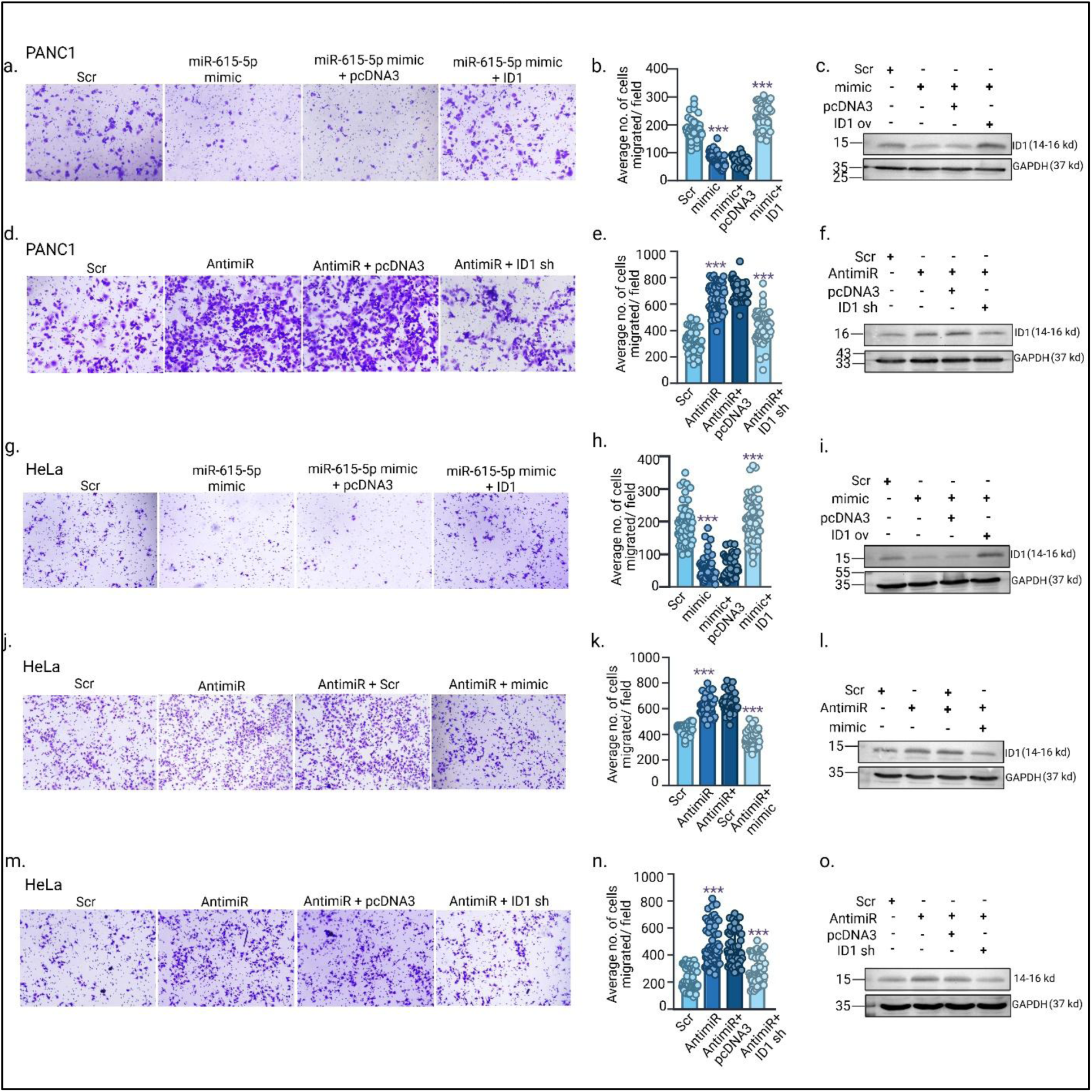
miR-615-5p suppresses ID1 mediated cellular migration. Role of miR-615-5p/ID1 interaction in the regulation of cancer cell migration was assessed by trans-well migration assay. a) The panel shows the representative microscopic fields of 3 independent trans-well migration assays, demonstrating the migratory potential of PANC1 cells under the influence of miR-615-5p overexpression and further ID1 complementation. b) Graphical representation of the average number of migrated cells per microscopic field in different experimental conditions are shown. For each experiment at least 10 microscopic fields were calculated for evaluating the average migrated cells per experimental conditions. The data is represented as average number of migrated cells ± SD of 3 independent experiments and significance was calculated using paired Students t-test (***p <0.0001). The number of cells migrated in each microscopic field was calculated using ImageJ software. c) Western immunoblotting shows the expression level of ID1 in mentioned experimental conditions. GAPDH was used as internal loading control. The image is a representative of 3 independent experiments. d) The figure shows the representative microscopic fields of 3 independent trans-well migration assays, demonstrating the migratory potential of PANC1 cells upon functional inhibition of miR-615-5p by AntimiR and followed by ID1 knockdown. e) Graphical representation of the average number of migrated cells per microscopic field in different experimental conditions are shown. For each experiment at least 10 microscopic fields were calculated for evaluating the average migrated cells per experimental conditions. The data is represented as average number of migrated cells ± SD of 3 independent experiments and significance was calculated using paired Students t-test (***p <0.0001). The number of cells migrated in each microscopic field was calculated using ImageJ software. f) Western immunoblotting shows the expression level of ID1 in mentioned experimental conditions. GAPDH was used as internal loading control. The image is a representative of 3 independent experiments. g) The panel represents the microscopic fields of 3 independent trans-well migration assays, demonstrating the migratory potential of HeLa cells under the influence of miR-615-5p overexpression which reverses upon further ID1 complementation. h) Graphical representation of the average number of migrated cells per microscopic field in different experimental conditions are shown. For each experiment at least 10 microscopic fields were calculated for evaluating the average migrated cells per experimental conditions. The data is represented as average number of migrated cells ± SD of 3 independent experiments and significance was calculated using paired Students t-test (***p <0.0001). i) Western immunoblotting shows the expression level of ID1 in mentioned experimental conditions. GAPDH was used as internal loading control. The image is a representative of 3 independent experiments. j) Specificity of miR-615-5p mediated ID1 regulation and its role in cellular migration was evaluated by a competition assay. The figure represents the microscopic fields of 2 independent trans-well migration assays, demonstrating the migratory potential of HeLa cells under functional inhibition of miR-615-5p followed by overexpression using mimic. k) Graphical representation of the average number of migrated cells per microscopic field in different experimental conditions are shown. For each experiment at least 10 microscopic fields were calculated for evaluating the average migrated cells per experimental conditions. The data is represented as average number of migrated cells ± SD of 2 independent experiments and significance was calculated using paired Students t-test (***p <0.0001). l). Western immunoblotting shows the expression level of ID1 in mentioned experimental conditions. GAPDH was used as internal loading control. The image is a representative of 3 independent experiments. m) The panel shows the representative microscopic fields of 3 independent trans-well migration assays, demonstrating the migratory potential of HeLa cells upon functional inhibition of miR-615-5p by AntimiR and followed by ID1 knockdown. n) Graphical representation of the average number of migrated cells per microscopic field in different experimental conditions are shown. For each experiment at least 15 microscopic fields were calculated for evaluating the average migrated cells per experimental conditions. The data is represented as average number of migrated cells ± SD of 3 independent experiments and significance was calculated using paired Students t-test (***p <0.0001). The number of cells migrated in each microscopic field was calculated using ImageJ software. o) Western immunoblotting shows the expression level of ID1 in mentioned experimental conditions. GAPDH was used as internal loading control. The image is a representative of 3 independent experiments.

### miR-615-5p suppresses ID1 mediated autophagy to regulate cellular migration

Autophagy is an evolutionarily conserved cellular process that primarily functions to maintain intracellular homeostasis through the recycling of damaged organelles and macromolecules under physiological stress. In cancer, however, this adaptive mechanism is frequently co-opted to support tumour cell survival, metabolic flexibility, and disease progression (27). Among solid malignancies, pancreatic ductal adenocarcinoma (PDAC) is particularly notable for its constitutively elevated autophagic activity and pronounced dependence on autophagy during tumour progression (25). Studies have shown that biallelic deletion of key autophagy regulators such as ATG5 or ATG7 markedly suppresses late-stage disease progression and the transition from pancreatic intraepithelial neoplasia (PanIN) to PDAC, while exerting comparatively limited effects on earlier disease initiation events such as acinar-to-ductal metaplasia (ADM) and early PanIN formation (42, 43). This stage-selective requirement is consistent with clinical and experimental observations of increased LC3B-II expression in advanced PDAC clinical samples (44). Recent reports have also implicated the role of ID1 in the regulation of autophagic responses in ovarian cancer cells, where it contributes to adaptive survival and therapy resistance. In order to explore a broader context in which ID1 and its regulation by miR-615-5 may intersect with autophagy and associated tumour phenotypes in PDAC, we examined whether miR-615-5p influences cellular autophagy. PANC1 and HeLa cells were transfected with a miR-615-5p mimic **(Figure 6a-b)** or a sequence-specific AntimiR **(Figure 6c-d)**. Ectopic over-expression of miR-615-5p resulted in a marked reduction in LC3B-II levels in both PANC1 **(Figure 6a, Supplementary fig. S5a** ∼ 50%, p< 0.0001, n=3) and HeLa **(Figure 6b, Supplementary fig. S5c** ∼ 50%, p= 0.0004, n= 3) cells, accompanied by a concomitant decrease in ID1 expression (PANC1= ∼ 50%, p< 0.0001, n=3; HeLa= ∼ 60%, p< 0.05, n=3) **(Figure 6a-b; Supplementary fig. S5b, Supplementary fig. S5d**). Conversely, functional inhibition of miR-615-5p using AntimiR led to an increase in LC3B-II abundance relative to scrambled and non-transfected controls in both PANC1 **(Figure 6c, Supplementary figure 5e,** fold change= ∼ 3-fold, p < 0.05, n= 3) as well as in HeLa **(Figure 6d, Supplementary fig. S5g,** fold change= ∼ 2-fold, p < 0.0001, n= 5) cells. AntimiR-mediated increase in autophagic response as indicated by increased expression of LC3B-II was accompanied with a concomitant elevation of ID1 expression (PANC1= ∼ 4-fold, p< 0.005, n=3 and HeLa= ∼ 2-fold, p <0.0001, n=5) (**Figure 6c, Supplementary fig. S5f, Figure 6c, Supplementary fig. S5h)** indicating that modulation of miR-615-5p reciprocally regulates both ID1 expression as well as LC3B-II abundance. This further suggests that miR-615-5p-mediated regulation of autophagy might be ID1-dependent. To determine whether the observed changes in LC3B-II reflect alterations in autophagic flux, cells were subjected to autophagic turnover assays using chloroquine (CQ), a late-stage autophagy inhibitor. Following 46h of microRNA modulation, cells were treated with CQ for 2h, and LC3B-II accumulation was assessed. Inhibition of miR-615-5p resulted in elevated autophagic flux in both PANC1 and HeLa cells; (Increased flux: PANC1= ∼ 2.6-fold, p= 0.0006, n=2; HeLa= ∼ 1.67-fold, p <0.005, n=3). In contrast, miR-615-5p overexpression led to reduced LC3B-II levels, and overall flux in both PANC1 **(Figure 6e-f, Supplementary fig. S5i-j)** and HeLa **(Figure 6g-h, Supplementary fig. S5k-l)** cells compared to control cells (PANC1= ∼ 80%, n=2, p <0.005; HeLa= 70%, n=3, p <0.005). Together, these findings suggest that miR-615-5p modulates autophagic flux in these cell models.

**Figure 6:**
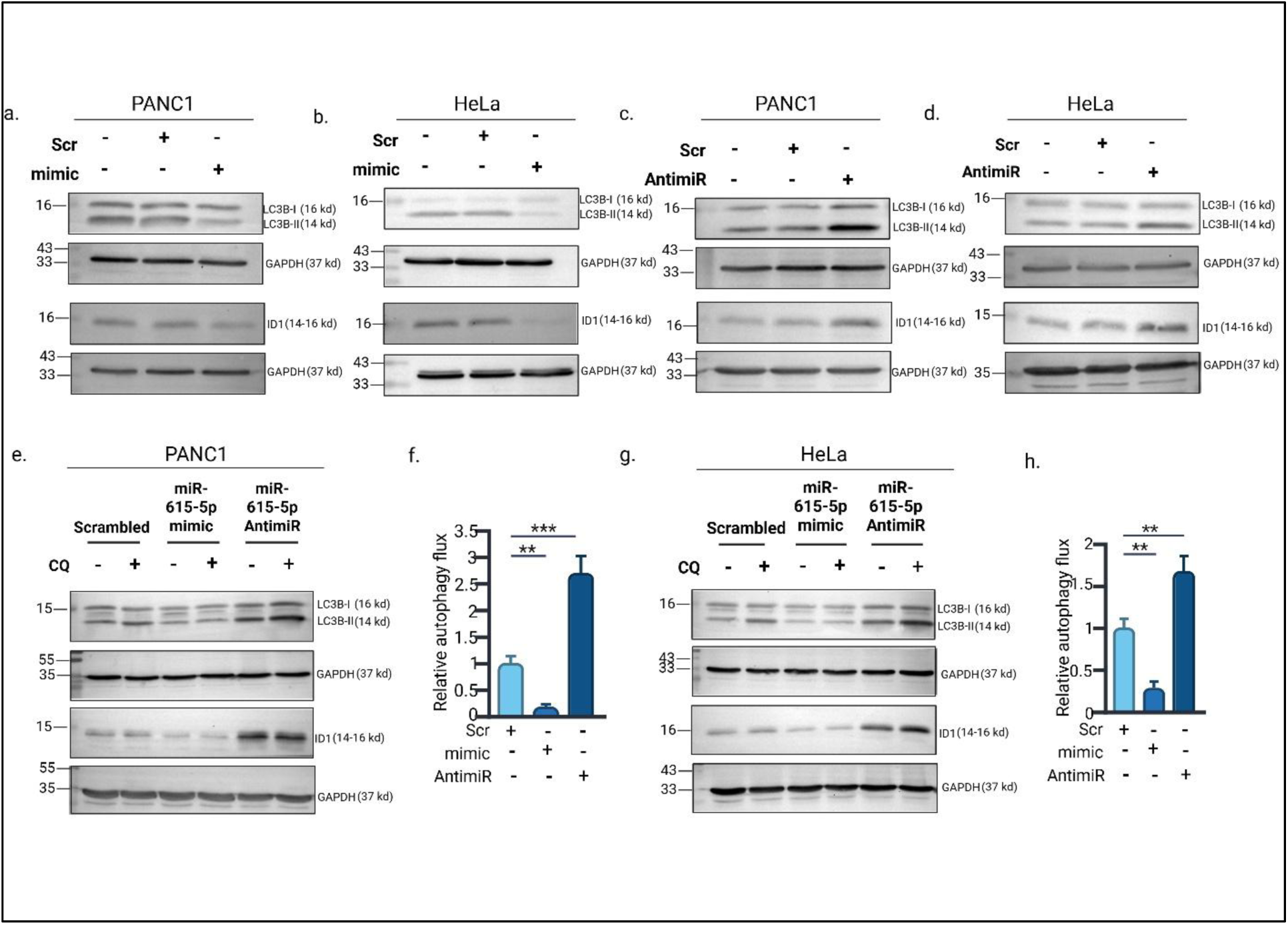
miR-615-5p suppresses cellular autophagy. Role of miR-615-5p in regulating cellular autophagy was assessed by evaluating LC3B-II expression in miR-615-5p gain of function and loss of functions system in PANC1 and HeLa cells a) Western-immunoblotting image showing the expression of LC3B-I/LC3B-II and ID1 protein expression upon miR-615-5p overexpressed (20nM) condition in PANC1 cells. GAPDH was used as internal loading control for both LC3B and ID1. The image is representative of 3 independent experiments. b) Western-immunoblotting image showing the expression of LC3B-I/LC3B-II and ID1 protein expression upon miR-615-5p overexpressed (15nM) condition in HeLa cells. GAPDH was used as internal loading control for both LC3B and ID1. The image is representative of 3 independent experiments. c) The figure represents the western immunoblotting image demonstrating the expression of LC3B-I/LC3B-II and ID1 protein expression upon inhibition of miR-615-5p in PANC1 cells by sequence specific AntimiR (25nM). GAPDH was used as internal loading control for both LC3B and ID1. The image is representative of 3 independent experiments. d) The figure represents the western immunoblotting image demonstrating the expression of LC3B-I/LC3B-II and ID1 protein expression upon inhibition of miR-615-5p in HeLa cells by sequence specific AntimiR (25nM). GAPDH was used as an internal loading control for both LC3B and ID1. The image is representative of 3 independent experiments. e) To evaluate autophagy flux, cells were treated with late-stage autophagy blocker CQ (25µM) for 2 hours. The LC3B-II expression in the untreated cells was subtracted from CQ treated cells (LC3B-II^CQ Treated^ – LC3B-II^CQ Untreated^). The net LC3B-II expression implies autophagy flux, which was then compared among different experimental conditions to assess the role of miR-615-5p in regulating autophagy flux. The figure represents western immunoblotting image of LC3B-I/LC3B-II and ID1 expression upon miR-615-5p overexpressed and inhibited condition in the presence and absence of CQ treatment in PANC1 cells. GAPDH was used as internal loading control for both LC3B and ID1. The image is representative of 2 independent experiments. f) Bar graph representing the autophagy flux in different experimental conditions and normalized against scrambled (Scr) control. The autophagy flux calculated from miR-615-5p mimic (LC3B-II^CQ Treated^ – LC3B-II^CQ Untreated^) and AntimiR (LC3B-II^CQ Treated^ – LC3B-II^CQ Untreated^) treated cells was normalized with scrambled (Scr) control (LC3B-II^CQ Treated^ – LC3B-II^CQ Untreated^) and represented graphically. The result shows that overexpression of miR-615-5p suppresses the autophagic flux by more than 80% whereas functional miR-615-5p inhibition upregulates it by 2.6-fold in PANC1 cells. The data is represented as the normalized average autophagy flux ± SE and significance was calculated using two-tailed paired Students t-test (**p <0.001; ***p <0.0001). g) The figure represents western immunoblotting image of LC3B-I/LC3B-II and ID1 expression upon miR-615-5p overexpressed and inhibited condition in the presence and absence of CQ treatment in HeLa cells. GAPDH was used as internal loading control for both LC3B and ID1. The image is representative of 3 independent experiments. h) Bar graph representing the autophagy flux in different experimental conditions and normalized against scrambled (Scr) control in HeLa cells. The autophagy flux calculated from miR-615-5p mimic (LC3B-II^CQ Treated^ – LC3B-II^CQ Untreated^) and AntimiR (LC3B-II^CQ Treated^ – LC3B-II^CQ Untreated^) treated cells was normalized with scrambled (Scr) control (LC3B-II^CQ Treated^ – LC3B-II^CQ Untreated^) and represented graphically. The result shows that overexpression of miR-615-5p suppresses the autophagic flux by ∼70% whereas inhibition upregulates it by 1.6-fold in HeLa cells. The data is represented as the normalized average autophagy flux ± SE and significance was calculated using two-tailed paired Students t-test (**p <0.001).

### miR-615-5p-mediated regulation of autophagy is ID1-dependent

Given the positive association observed in ID1 expression as well as LC3B-II levels, the contribution of ID1 to miR-615-5p mediated regulation of autophagy was further investigated. Inhibition of miR-615-5p using AntimiR is accompanied by an increased LC3B-II expression in both PANC1 **(Figure 7a, Supplementary fig. S6a)** as well as in HeLa **(Figure 7b, Supplementary fig. S6c)** cells, indicating upregulation of autophagy. shRNA mediated silencing of ID1 in both PANC1 and HeLa cells **(Figure 7a-b)** caused significant decrease in the expression of LC3B-II in PANC1 **(Figure 7a**, **Supplementary fig. S6a,** decrease = ∼ 50 %, p < 0.0001, n= 2) and HeLa **(Figure 7b**, **Supplementary fig. S6c,** decrease = ∼ 30%, p< 0.005, n= 3) cells, thereby indicating that miR-615-5p-mediated regulation of autophagy is ID1-dependent. To further confirm this, autophagy was assessed upon restoration of ID1 expression in mimic transfected HeLa cells. In miR-615-5p overexpressing HeLa cells, reintroduction of ID1 restored LC3B-II levels relative to miR-615-5p mimic treated cells (**Figure 7c**, **Supplementary fig. S6e,** fold change = ∼ 2.1-fold, n= 4, p <0.005), confirming that changes in the LC3B-II levels following miR-615-5p modulation are dependent, at least in part, on ID1 expression. Previous experiments established a role for the miR-615-5p/ID1 axis in regulating cellular migration *in vitro* **(Figure 5)**. To assess whether miR-615-5p-driven autophagic induction also promotes migration, PANC1 cells were pretreated with CQ to impair autophagic progression prior to migration assays. While ectopic expression of ID1 rescued the mimic-mediated inhibition of cellular migration in miR-615-5p overexpressing cells, CQ-mediated inhibition of autophagy resulted in significantly decreased migration (∼ 50%, n=3, p <0.0001) of PANC1 cells (**Figure 7d-e**) even under ID1 over-expression, thereby indicating that miR-615-5p/ID1-axis driven migration is autophagy-dependent. The outcome of these experiments identified a crucial involvement of ID1 mediated autophagy in the suppression of cellular migration by miR-615-5p. Collectively, these findings identify an miR-615-5p/ID1 regulatory axis in PDAC cells and demonstrate that miR-615-5p suppresses ID1 expression, with downstream effects on autophagy-associated migratory behaviour. These data define a previously unrecognized regulation of ID1 and support a functional link between miR-615-5p suppression, ID1 overexpression, and its associated pro-migratory phenotypes in PDAC.

**Figure 7:**
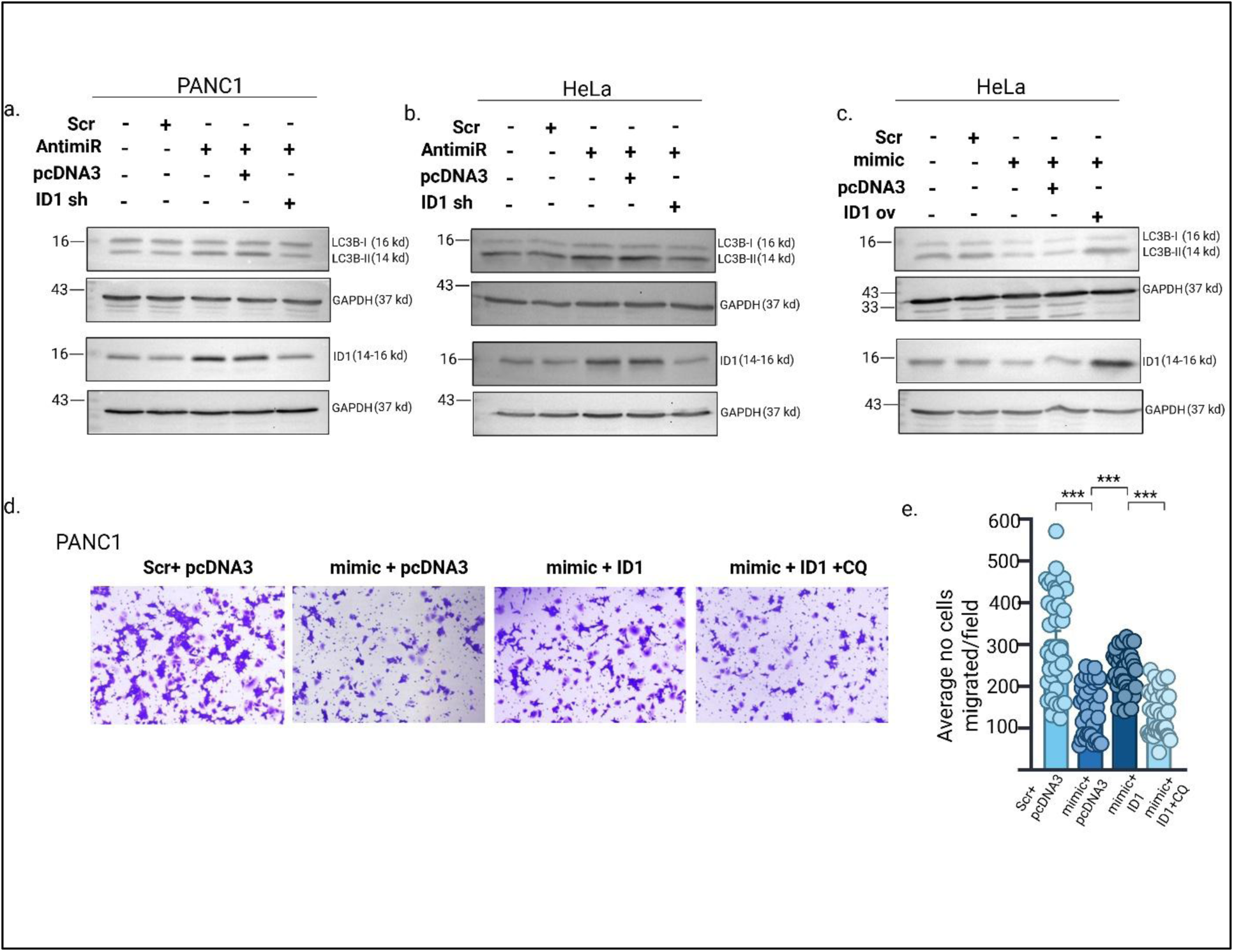
miR-615-5p suppresses ID1 mediated autophagy to regulate cellular migration. Role of ID1 in miR-615-5p mediated regulation of autophagy was assessed either by overexpressing ID1 in the presence of mimic or suppressing ID1 upon the functional inhibition of miR-615-5p. LC3B-II and ID1 expression was assessed in these experimental conditions in both PANC1, and HeLa cells. a) The western-immunoblot result demonstrated the expression of LC3B-I/LC3B-II and ID1 expression upon the inhibition of miR-615-5p by transfecting miR specific AntimiR (25nM) followed by shRNA (1µg) mediated knockdown of ID1 in PANC1 cells. GAPDH is used as internal loading control. The image is the representation of 2 independent experiments. b) The western-immunoblot result demonstrated the expression of LC3B-I/LC3B-II and ID1 expression upon the inhibition of miR-615-5p by transfecting miR specific AntimiR (25nM) followed by shRNA (1µg) mediated knockdown of ID1 in HeLa cells. GAPDH is used as internal loading control. The image is the representation of 3 independent experiments. c) The result demonstrated the expression of LC3B-I/LC3B-II and ID1 after ectopic overexpression of miR-615-5p (15nM) followed by the complementation of ID1 (500ng) in HeLa. GAPDH is used as internal loading control for both LC3B-II and ID1. The image is the representation of 4 independent experiments. d) The role of autophagy in miR-615-5p/ID1 mediated cellular migration was evaluated by assessing the cellular migratory potential by chemically inhibiting cellular autophagy by treating the cells with CQ (25µM) 3 hours prior to trans-well migration. The panel shows the representative microscopic fields of 3 independent trans-well migration assays, demonstrating the migratory potential of PANC1 under different experimental conditions. e) The bar graph represented the average number of migrated cells per microscopic field in different experimental conditions. For each experiment at least 15 microscopic fields were calculated for evaluating the average migrated cells per experimental conditions. The data is represented as average number of migrated cells ± SD of 3 independent experiments and significance was calculated using paired Students t-test (***p <0.0001). The number of cells migrated in each microscopic field was calculated using ImageJ software.

## Discussion

In PDAC, high expression of ID1 is often associated with enhanced metastasis, uncontrolled proliferation, therapeutic resistance leading to accelerated tumorigenesis which established ID1 as one of the crucial oncogenes in PDAC (10, 11). Considering its consistent overexpression and associated tumorigenic activities, Tang et. al. suggested ID1 as a potential biomarker for PDAC (11). So far, only miR-29b and miR-381 were identified as direct regulators of ID1 which function through target-specific interactions in human lung adenocarcinoma (17–19), although reports of miR-mediated post transcriptional regulation of ID1 and its consequence in tumorigenesis is relatively scarce. In the present study, we identify and functionally validate a previously uncharacterized regulatory axis in PDAC where tumour-suppressor miR-615-5p directly targets the 3′-UTR of the oncogene ID1 and modulates ID1 dependent cellular migration through regulating cellular autophagy. These findings extend our current understanding of ID1 regulation by establishing a microRNA-mediated layer of control with mechanistic and functional consequences in the pathogenesis of PDAC.

A study by Trevino et. al. (44) studied the expression pattern of ID1 gene across the different grades of PDAC and how ID1 expression affects the prognostic outcome of the disease. To analyse the expression of ID1, total 140 PDAC patients were evaluated of which 8 grade I, 83 grade II, 48 grade III, and 1 grade IV patients were included. A significant positive correlation (Spearmen’s correlation: 0.29, p-value = 0.0006) was observed in ID1 expression with the subsequent progression of the disease (10). High expression of ID1 in these cells enables them to escape TGF-beta mediated apoptosis leading to tumour progression, highlighting a crucial role of ID1 expression in PDAC progression. PDAC is likewise associated with reduced expression of miR-615-5p, which has been categorized as a tumour-suppressive microRNA due to its ability to inhibit cellular migration and invasion *in vitro* and to restrain tumour growth *in vivo*. In this study, the *in silico* screening predicted miR-615-5p as a putative interactor of ID1 followed by analysis of the PDAC clinical tissue samples which highlighted a significant inverse correlation between miR-615-5p and ID1 (**Figure 1f-i**). Hence, the *in silico* analysis, along with prior evidence supporting the oncogenic role of ID1 and the tumour-suppressive function of miR-615-5p in PDAC, prompted us in further investigating a potential regulatory relationship between these two molecules.

Identification of miRNA regulators of oncogenic ID1 was guided by cross-platform *in silico* prediction using TargetScan, miRwalk and miRSystem (**Figure 1f**). Among the commonly predicted miRNAs, miR-615-5p was selected based on its predicted binding stability and seed complementarity within the ID1 3′-UTR **(Figure 1g)**, together with prior reports describing its tumour-suppressive role in PDAC. Moreover, among the predicted miRs, miR-615-5p showed a significant inverse correlation with ID1 expression in PDAC clinical samples **(Figure 1i)**. While additional predicted miRNAs were not functionally examined in the present study, it is well established that individual transcripts are frequently regulated by multiple miRNAs acting in combinatorial networks (40). Therefore, miR-615-5p mediated regulation likely represents one component of a broader post-transcriptional landscape governing ID1 expression using a pan analysis of micro RNAs regulating ID1 expression, which warrants further systematic investigation. Expression analysis of ID1 upon miR-615-5p overexpression and functional inhibition showed a significant downregulation and upregulation of ID1 **(Figure 2a-e)**, respectively. Moreover, reintroduction of miR-615-5p in the presence of AntimiR reversed the expression of ID1, highlighting the specificity of miR activity on ID1 gene expression **(Figure 2i-j)**. Luciferase reporter assays designed to validate the predicted interaction between miR-615-5p and the ID1 3′-UTR showed that miR-615-5p suppresses ID1 3′-UTR activity in a sequence-specific manner, as disruption of the predicted binding site failed to repress the luciferase activity **(Figure 3b),** thereby justifying the hypothesis of miR-615-5p-ID1 inhibitory interaction. Notably, mutation of the miR-615-5p binding site did not increase basal reporter activity relative to the wild-type construct. This observation suggests the presence of endogenous regulatory mechanisms that continue to constrain ID1 3′-UTR activity despite loss of the miR-615-5p site. Because gene expression is frequently regulated by multiple miRNAs acting in combination, disruption of a single miRNA binding site would be expected to selectively eliminate miR-615-5p mediated repression while leaving other regulators intact. Such buffering by parallel miRNA regulators provides a plausible explanation for the preserved basal activity and highlights the layered nature of post-transcriptional gene control. Furthermore, the interaction between miR-615-5p and ID1 3’-UTR was substantiated by several biochemical enrichment assays using RIP. From biotinylated RIP assays, it was demonstrated that miR-615-5p directly associates with the 3’-UTR of ID1 gene which supports the experimental outcome of luciferase reporter assay **(Figure 4a-b)**. *In vitro* AGO2 assay with gain of function and loss of function of miR-615-5p showed selective enrichment patterns of ID1 transcript which highlights its association with AGO2 and ID1 transcript and thereby identifying an underlying mechanism of ID1 regulation **(Figure 4e-f, 4i-j)**. Although AGO2 pulldown has been considered as a valuable biochemical tool for assessing the physical interaction between miRs with their targets however, it is often argued that experimental manipulation of miRNA levels through transfection may lead to broad transcriptomic alterations. These global changes can obscure the distinction between direct miRNA-mediated regulation and indirect downstream effects, thereby complicating the interpretation of AGO2-associated target enrichment. These unintentional, non-specific alterations may influence the association of miRNA with its target. To overcome this, the present study incorporated a RIP competition assay using AntimiR. Consistent with *in vitro* AGO2 RIP, the result from these AntimiR competition assays further confirmed the association of miR-615-5p with ID1 and AGO2 highlighting the specificity of this interaction as well as the formation of the functional RISC **(Figure 4m-n)**.

Metastasis is a multistep biological process involving local invasion, intravasation, survival in circulation, extravasation, and colonization at distant sites. Among these, cell migration and invasion are fundamental and rate-limiting early events that critically determine metastatic potential (45). *In vitro* migration assays provide a mechanistically relevant and experimentally tractable alternative to metastasis by enabling quantitative analysis of cellular motility, an essential determinant of metastatic potential under highly controlled conditions, thereby facilitating high-resolution mechanistic insight. Considering, miR-615-5p was earlier reported to suppress the migratory activity of the PDAC cells *in vitro* (21) and associated oncogenic activity of ID1 in highly metastatic PDAC (10, 11), following study focused on analysing the potential regulatory role of miR-615-5p/ID1 axis in modulating cellular migratory behaviour. Consistent with the previous reports, a reduction in the cellular migration upon miR-615-5p overexpression was also observed in this study which reinforces the tumour-suppressive role of miR-615-5p (**Figure 5a-c** & **5g-i**). Further, complementation of ID1 in presence of miR-615-5p overexpression reverses the migratory potential of miR-615-5p overexpressed cells **(Figure 5a-c** & **5g-i**), thereby confirming that miR-615-5p-mediated regulation of cell migration is ID1-dependent. This observation was further validated, where induction in migration owing to functional inhibition of miR-615-5p was significantly reduced upon ID1 knock-down highlighting the role of ID1 in miR-615-5p mediated regulation of cellular migration **(Figure 5d-f & 5m-o)**. Additionally, the increased cellular migration upon functional inhibition of miR-615-5p by AntimiR was also reversed by the introduction of miR-615-5p mimics, which is coupled with reciprocal decrease in ID1 expression level further substantiate the potential role of ID1 in miR-615-5p mediated regulation of cellular migration *in vitro* **(Figure 5j-l)**. However, miR-615-5p, being a tumour suppressor miRNA also regulates numerous target genes and most of which are unidentified. Thus, ID1 can be one of the many plausible targets of miR-615-5p to drive the suppression of cellular migration.

PDAC is characterized by high basal level autophagy and was monitored by LC3B-II level, which allows the lipidation and association with autophagic vesicles (25). As observed by Yang et. al., almost 81% of the high-grade PDAC clinical samples (65 of 80 PDAC) showed elevated LC3B-II expression compared to normal pancreatic ductal epithelium or low-grade pancreatic neoplasia (25). Elevated LC3B-II level was further reported to be associated with development and aggressiveness of the PDAC (42). These findings suggest an important role of autophagy in late-stage disease aggressiveness, including malignant growth and metastatic progression. Recent study by Meng et. al. highlighted the potential role of ID1 in ovarian cancer, where elevated expression of ID1 is associated with increased autophagy and associated therapeutic resistance (28). Considering the high expression levels of ID1 in PDAC and its reported association with autophagy coupled with the suppressive nature of miR-615-5p, we extended our study to assess the role of miR-615-5p/ID1 axis in regulating cellular autophagy. The outcome of the study showed that overexpression of miR-615-5p which downregulated ID1 expression also suppressed the basal LC3B-II expression, **(Figure 6a-b)** while autophagic flux assessment following CQ treatment shows a significant decrease in the autophagic turnover compared to the control cells **(Figure 6e-h)**. The reverse phenomenon was observed in presence of AntimiR **(Figure 6c-d & 6e-h)**. The outcome highlights a yet unexplored regulation of miR-615-5p on cellular autophagy and its plausible role in the pathogenesis of cancer. Involvement of ID1 in this regulation was also assessed, where complementation of ID1 in miR-615-5p overexpressed cells reversed LC3B-II expression **(Figure 7c)**, whereas, knocking down ID1 in presence of AntimiR significantly reduced the LC3B-II level as compared to AntimiR **(Figure 7a-b)**. Collectively, these results highlight the role of miR-615-5p/ID1 axis in regulating cellular autophagy. Previous results of this study pointed out a critical role of miR-615-5p in suppressing ID1 mediated cellular migration as an *in vitro* index of metastasis, where complementation of ID1 rescued the migratory potential of the miR-615-5p overexpressed cells **(Figure 5a & 5g)**. Interestingly, impairing autophagy by pre-treating the cells with CQ failed to achieve ID1 mediated cellular migration in PDAC cell line *in vitro,* pointing out a crucial involvement of ID1 mediated autophagy in the suppression of cellular migration by miR-615-5p **(Figure 7d-e)**.

Collectively, the experimental findings indicate that miR-615-5p acts as a negative regulator of ID1 expression and is associated with reduced cellular migration, at least in part through modulation of ID1-linked autophagic activity. The data support a regulatory relationship in which reduced miR-615-5p levels contribute to elevated ID1 expression in PDAC cellular models. This observation is consistent with prior reports linking higher ID1 expression with suppression of miR-615-5p leading to aggressive disease features in PDAC. Together, these results describe identification of a novel miR-615-5p/ID1 regulatory axis that adds a mechanistic framework to the upstream control of ID1 in this disease context.

## Data Availability

The data generated in this study are available within the article and its supplementary data files. All other raw data generated in this study are available upon request to the corresponding author. Publicly available expression datasets analyzed in this study were obtained from the Gene Expression Omnibus (GEO) under accession numbers GSE41368 and GSE41369. Additional publicly available datasets and resources used for expression and survival analyses are described in the Methods section.

## Ethics Statement

This study did not involve the collection or use of human subjects or human specimens by the authors. All human-derived clinical data analyzed in this study were obtained from publicly available datasets. Therefore, additional institutional ethical approval and informed consent were not required for this study.

## Authors’ Contributions

**AS:** Conceptualization, data curation, formal analysis, validation, investigation, visualization, methodology, writing–original draft, writing–review, editing and funding acquisition. **SR:** Data curation, formal analysis, investigation. **AR:** Data curation, software, formal analysis, investigation, visualization. **KB:** Conceptualization, resources, supervision, funding acquisition, project administration, validation, methodology, writing-original draft, writing-review and editing.

## Supporting information

Supplementary Figures with Legends

## Acknowledgments

This work is supported from intramural funding from Bose Institute, Kolkata, India, as well as from extramural funding from DST-SERB sponsored project CRG/2021/004623, dated 2022 and Indian Council of Medical Research (ICMR), New Delhi, Sanction number: 2019-0137-CMB/adhoc/BMS. A.S. acknowledges the University Grants Commission (U.G.C.) to provide his fellowship (341948). A.R. (201610053746) and S.R. (Nov2017-341475) acknowledges the University Grants Commission (U.G.C.) to provide their fellowship. We would like to thank all the staffs of Central Instrument Facility of Bose Institute and Mr. Mrinal Das for DNA sequencing. Generative AI tools were used to improve the clarity and readability of the manuscript and to assist in revising and translating author-written text for clearer communication. All scientific content was reviewed and verified by the authors.

## Note

Supplementary data for this article are available at Cancer Research Online (http://cancerres.aacrjournals.org/).

