## Supplementary Figures with Legends for "The miR-615-5p/ID1 Axis Regulates Autophagy-Associated Migration in Pancreatic Ductal Adenocarcinoma"

### Supplementary Figures and Legends:

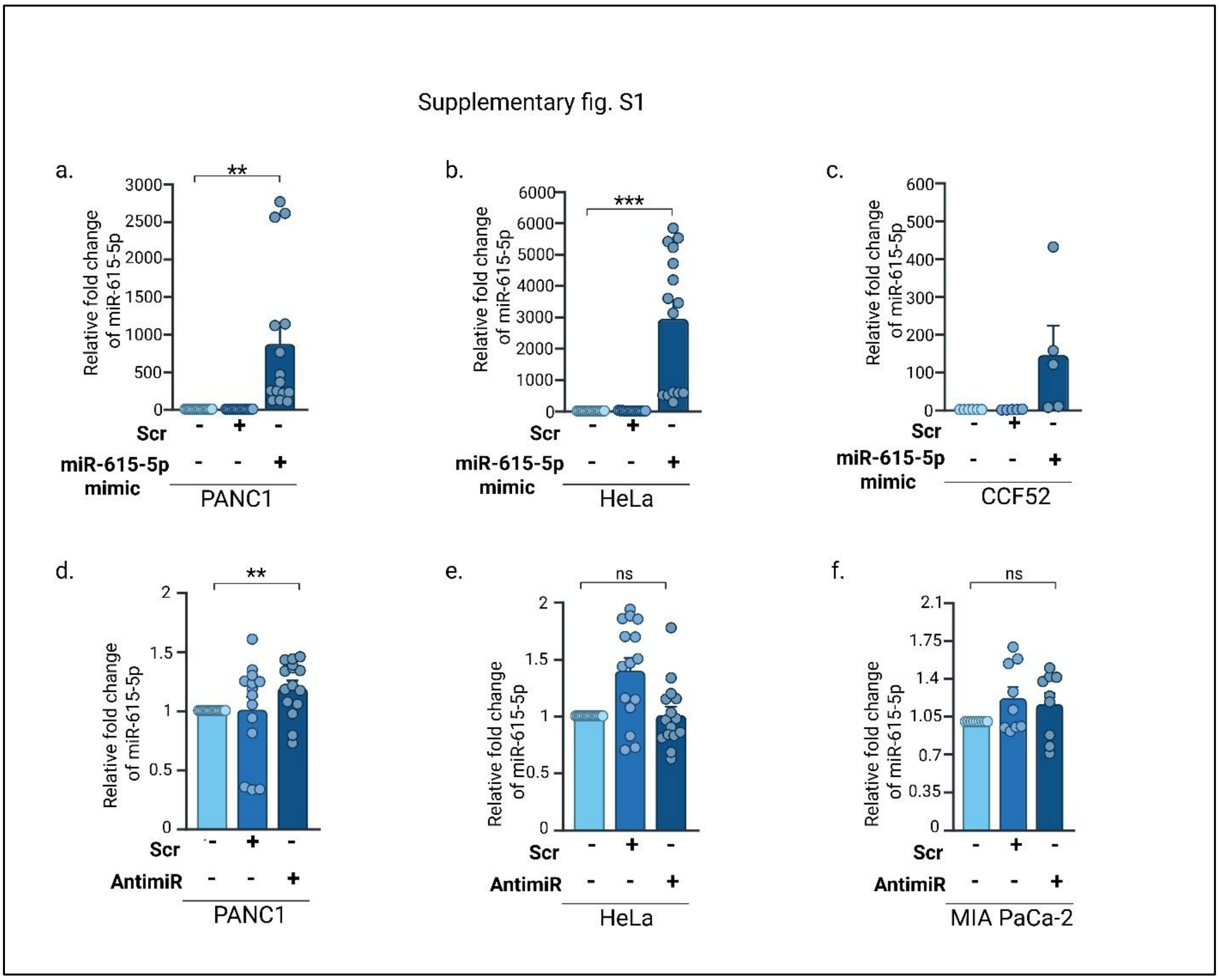


**Supplementary Figure S1: Expression analysis of miR-615-5p following mimic overexpression and functional inhibition (Corresponds to Figure 2).** miR-615-5p was ectopically overexpressed by transient transfection of a miR-615-5p-specific mimic, and its expression was assessed by real-time PCR (Fig: 2). (a) miR-615-5p expression in PANC1 cells following transfection with 20nM mimic. (b) miR-615-5p expression in HeLa cells following transfection with 15nM mimic. (c) miR-615-5p expression in CCF52 cells following transfection with 32.5nM mimic. Data are presented as mean ± SE from 5 independent experiments for PANC1 and HeLa cells and 2 independent experiments for CCF52 cells. Functional inhibition of miR-615-5p was achieved by transient transfection of a sequence-specific AntimiR (25nM), and miR-615-5p expression was subsequently assessed by real-time PCR. (d) miR-615-5p expression in PANC1 cells. (e) miR-615-5p expression in HeLa cells. (f) miR-615-5p expression in MIA PaCa-2 cells. Data are presented as mean ± SE from 5 independent experiments for PANC1 and HeLa cells and 5 independent experiments for MIA PaCa-2 cells. Statistical significance was determined using a paired Student's t-test (**p <0.001; ***p <0.0001).

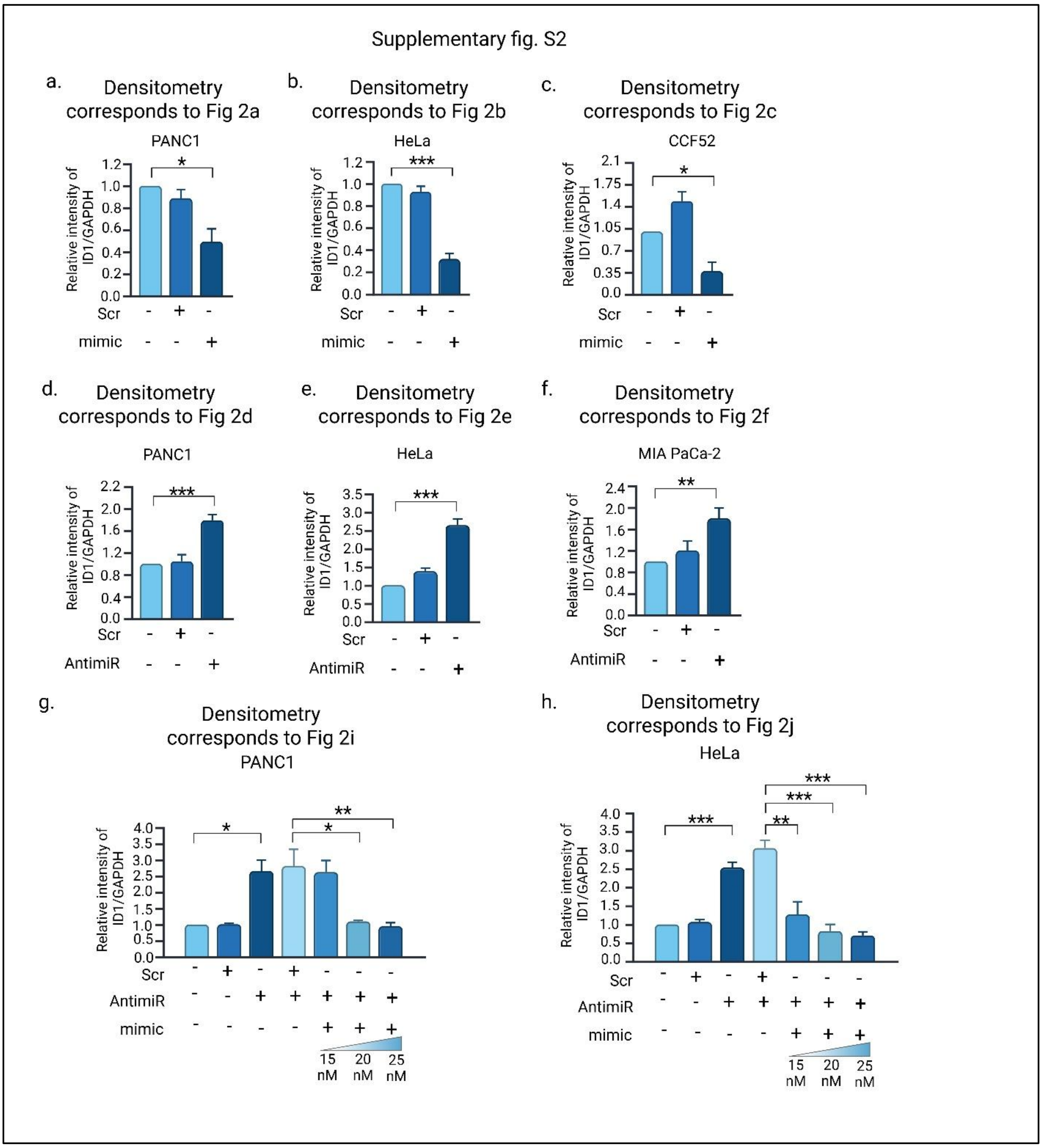


**Supplementary Figure S2: Densitometric quantification of ID1 protein expression following miR-615-5p mimic and AntimiR treatment (Corresponds to Figure 2).** miR-615-5p was ectopically overexpressed using a miR-615-5p-specific mimic or functionally inhibited using a sequence-specific antimiR, and ID1 protein expression was evaluated by western immunoblotting (Figure 2). (a) ID1 expression in PANC1 cells following transfection with 20nM miR-615-5p mimic, corresponding to Figure 2a. (b) ID1 expression in HeLa cells following transfection with 15nM miR-615-5p mimic, corresponding to Figure 2b. (c) ID1 expression in CCF52 cells following transfection with 32.5nM miR-615-5p mimic, corresponding to Figure 2c. (d) ID1 expression in PANC1 cells following transfection with 25nM antimiR, corresponding to Figure 2d. (e) ID1 expression in HeLa cells following transfection with 25nM antimiR, corresponding to Figure 2e. (f) ID1 expression in MIA PaCa-2 cells following transfection with 25nM antimiR, corresponding to Figure 2f. (g-h) ID1 expression in PANC1 and HeLa cells following antimiR-

mediated inhibition of miR-615-5p (25nM) and subsequent reintroduction of miR-615-5p mimic at increasing concentrations (15, 20, and 25nM), respectively. GAPDH was used as the loading control for PANC1, HeLa, and MIA PaCa-2 cells, whereas β-actin was used for CCF52 cells. Densitometric analysis was performed using ImageJ. ID1 protein levels were normalized to the corresponding loading control, and relative protein expression was calculated with respect to the respective control, which was set to 1. Data are presented as mean ± SE from at least 3 independent experiments for PANC1, HeLa, and MIA PaCa-2 cells and 2
independent experiments for CCF52 cells. For figure c, the significance was calculated using two-tailed paired Students t-test using 2 different exposures of each experiment. Statistical significance was determined using a paired two-tailed Student's t-test (*p <0.05; **p <0.001; ***p <0.0001).

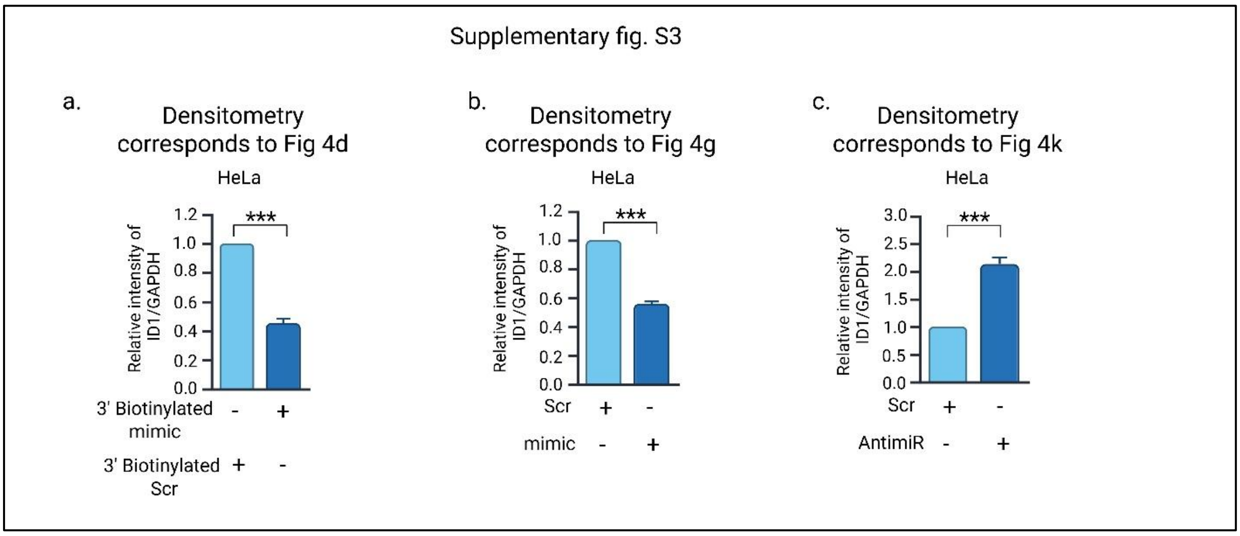


**Supplementary Figure S3: Densitometric quantification of ID1 protein expression following miR-615-5p modulation before RIP analysis (Corresponds to Figure 4).** a) 3'Biotinylated miR- 615-5p (15nM) was ectopically overexpressed in HeLa cells and its association with ID1 was assessed by RIP assay. ID1 protein expression was evaluated by western immunoblotting upon 3'Biotinylated miR-615-5p overexpression. b) miR-615-5p was overexpressed using a miR-615-5p-specific mimic (15nM). ID1 expression was evaluated before evaluating its association with AGO2- RISC complex using RIP assay in HeLa cells. c) miR-615-5p was functionally inhibited using a sequence-specific antimiR (25nM) to evaluate its association with AGO2-RISC complex using RIP assay in HeLa cells. ID1 protein expression was evaluated by western immunoblotting in these experimental conditions. GAPDH was used as the loading control. Densitometric analysis of western blot bands was performed using ImageJ software. ID1 protein levels were normalized to GAPDH and relative protein expression was calculated with respect to control samples, which were set to 1. Data are presented as mean ± SE from at least three independent experiments. Statistical significance was determined using paired two-tailed Student's t-test. (***p < 0.001).

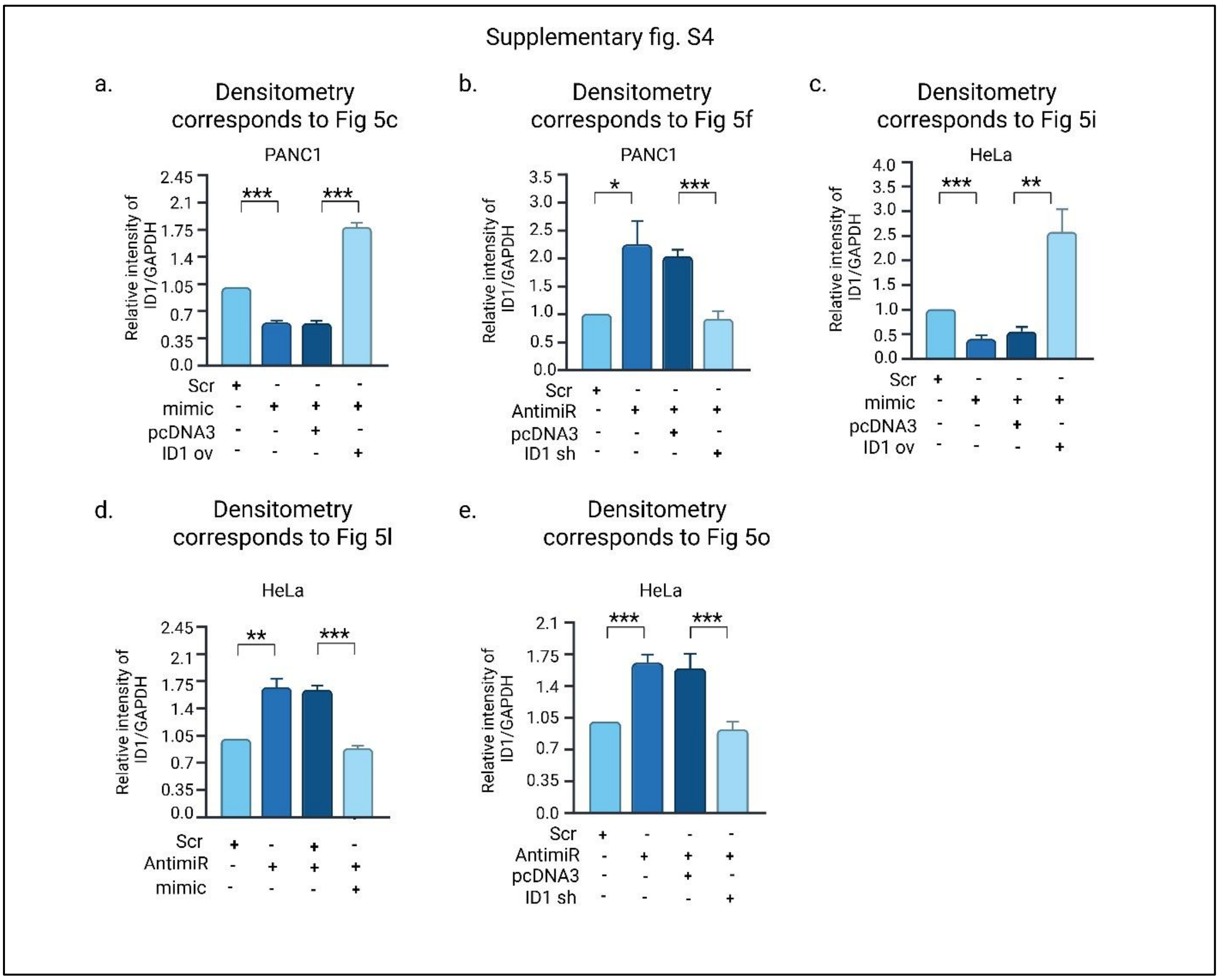


**Supplementary Figure S4: Densitometric analysis of ID1 expression to assess miR-615-5p/ID1 interaction in the regulation of cancer cell migration (Corresponds to Figure 5).** a) Densitometric analysis of ID1 in the mentioned experimental conditions in PANC1 cells. b) Densitometric analysis of ID1 in the mentioned experimental conditions in PANC1 cells. c) Densitometric analysis of ID1 in the mentioned experimental conditions in HeLa cells. d) Densitometric analysis of ID1 in the mentioned experimental conditions in HeLa cells. e) Densitometric analysis of ID1 in the mentioned experimental conditions in HeLa cells. GAPDH was used as the internal loading control for all the experiments. Densitometric analysis of western blot bands was performed using ImageJ software. ID1 protein levels were normalized to GAPDH and relative protein expression was calculated with respect to control samples, which were set to 1. Data are presented as mean ± SE from at least three independent experiments. Statistical significance was determined using paired two-tailed Student's t-test (*p <0.05; **p <0.001; ***p <0.0001).

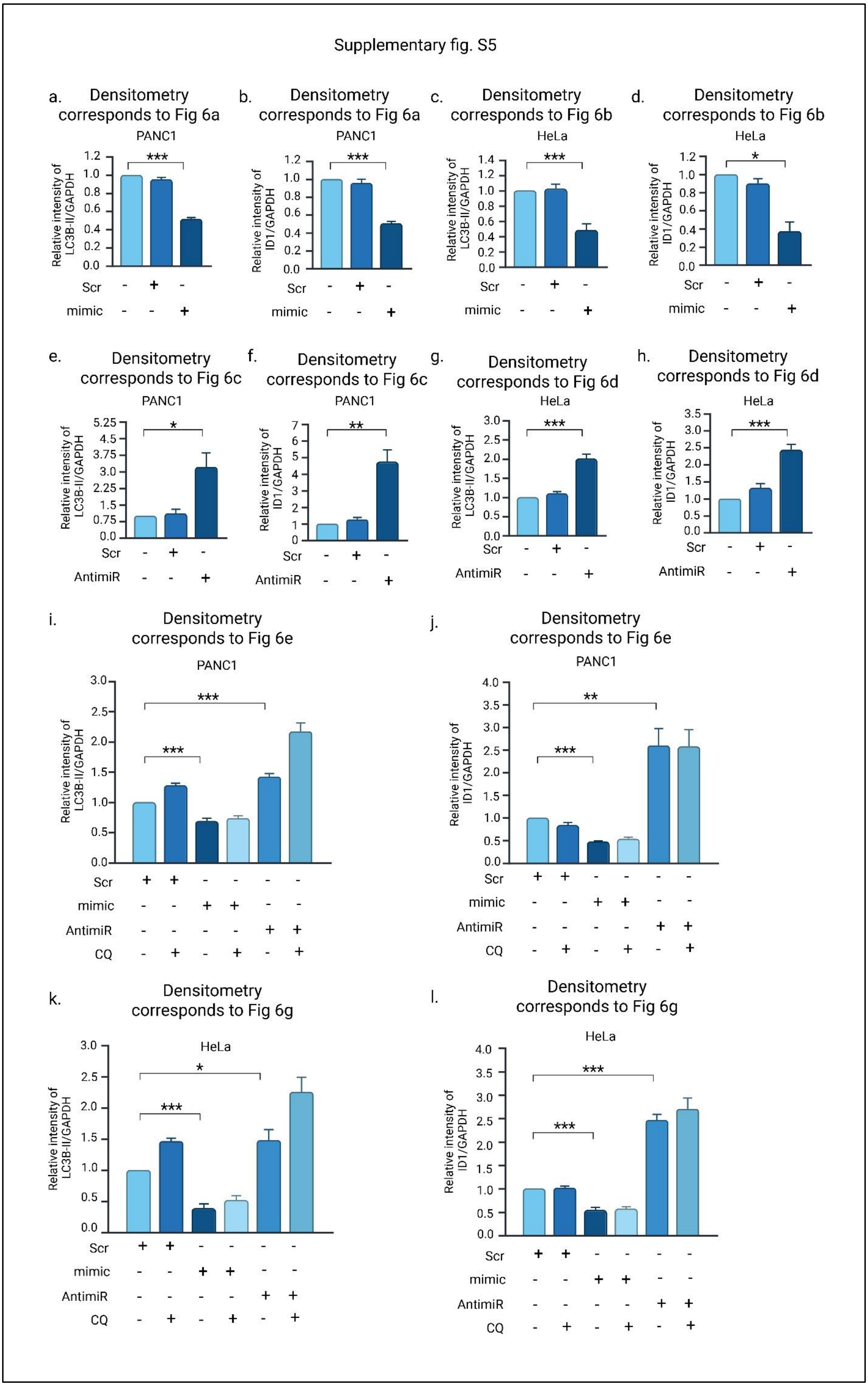


**Supplementary Figure S5: Densitometric analysis of ID1 expression to evaluate the role of**

**miR- 615-5p/ID1 regulation on cellular autophagy (Corresponds to Figure 6).** a-b) Densitometric analysis of ID1 and LC3B-II upon 20nM miR-615-5p mimic overexpressed condition in PANC1

cells, respectively. c-d) Densitometric analysis of ID1 and LC3B-II upon 15nM miR-615-5p mimic overexpressed condition in HeLa cells, respectively. e-f) Densitometric analysis of ID1 and LC3B-II upon functional inhibition of miR-615-5p by AntimiR (25nM) PANC1 cells, respectively. g-h) Densitometric analysis of ID1 and LC3B-II upon functional inhibition of miR-615-5p by AntimiR (25nM) HeLa cells, respectively. i-j) Densitometric analysis of ID1 and LC3B-II in the mentioned experimental conditions in PANC1 cells, respectively. k-l) Densitometric analysis of ID1 and LC3B-II in the mentioned experimental conditions in HeLa cells, respectively. GAPDH was used as the internal loading control for all the experiments. Densitometric analysis of western blot bands was performed using ImageJ software. ID1 protein levels were normalized to GAPDH and relative protein expression was calculated with respect to control samples, which were set to 1. Data are represented as mean ± SE from at least 3 independent experiments, whereas k-l is represented as mean ± SE of 2 independent experiments. The significance was calculated using two-tailed paired Students t-test using 3 different exposures of each experiment. For the other experiments the statistical significance was determined using paired two-tailed Student's t-test. (*p <0.05; **p <0.001; ***p <0.0001).

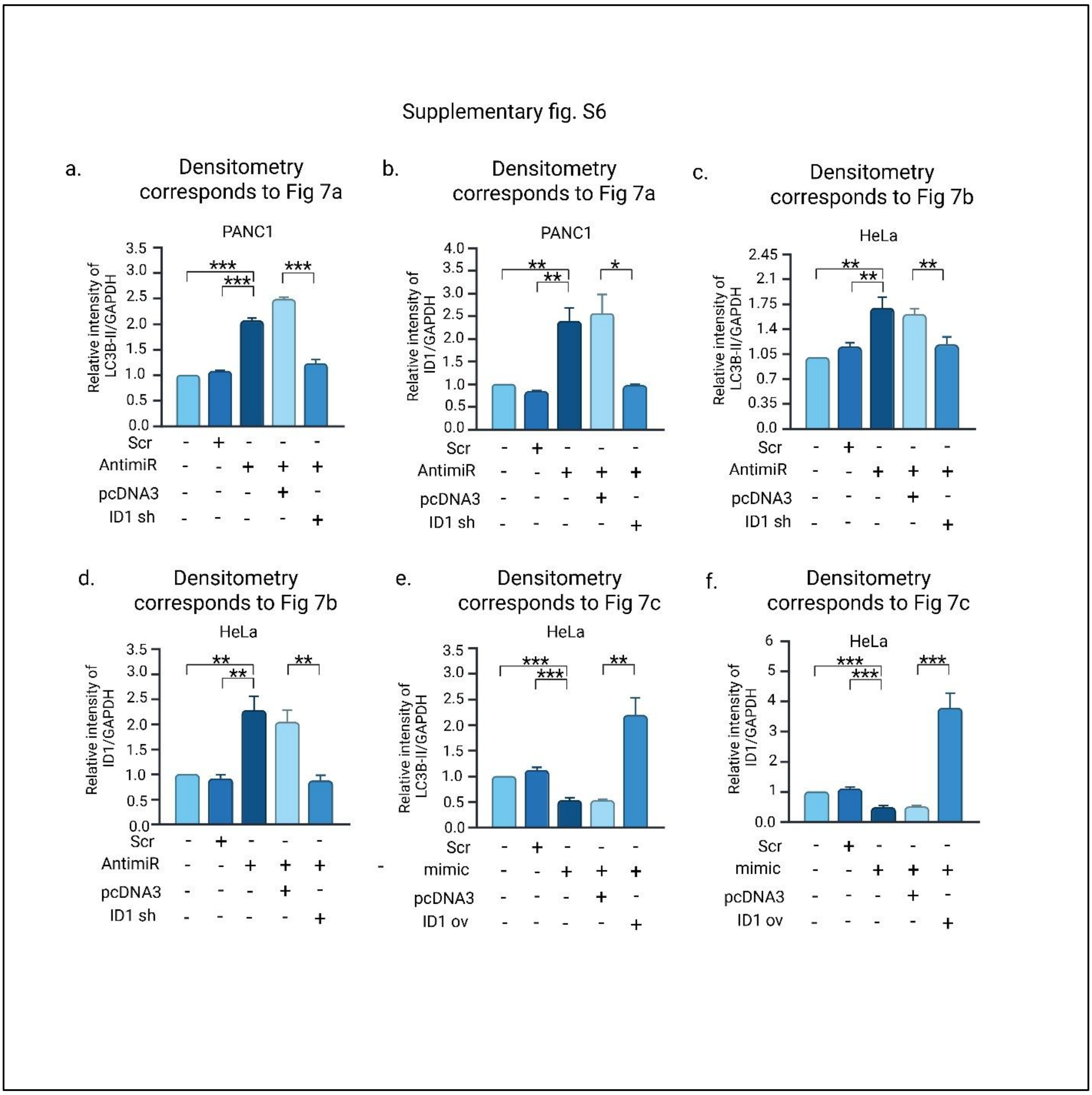


**Supplementary Figure S6: Densitometric analysis of ID1 expression to assess the role of ID1 in miR-615-5p mediated regulation of cellular autophagy (Corresponds to Figure 7).** a-b) Densitometric analysis of ID1 and LC3B-II in the mentioned experimental conditions in PANC1 cells, respectively. c-d) Densitometric analysis of ID1 and LC3B-II in the mentioned experimental conditions in HeLa cells, respectively. e-f) Densitometric analysis of ID1 and LC3B-II in the mentioned experimental conditions in HeLa cells, respectively. GAPDH was used as the internal loading control for all the experiments. Densitometric analysis of western blot bands was performed using ImageJ software. ID1 protein levels were normalized to GAPDH and relative protein expression was calculated with respect to control samples, which were set to 1. Data are presented as mean ± SE from at least 3 independent experiments, whereas a-b is represented as mean ± SE of 2 independent experiments. The significance was calculated using two-tailed paired Students t-test using 3 different exposures of each experiment. For other experiments statistical significance was determined using paired two-tailed Student's t-test. (*p <0.05; **p <0.001; ***p <0.0001).

**Supplementary Table S7:**
**List of the primers used in this study**

| | |
|---|---|
| ID1 Forward primer | 5' TCCTCTCTGCACACCTACTA 3' |
| ID1 Reverse primer | 5' GCACCAAACGTGACCATTT 3' |
| hsa-miR-615-5p forward primer | 5' GGGGGTCCCCGGTGCTCGGATC 3' |
| Poly(T) adaptor | 5' GCGAGCACAGAATTAATACGACTCACTATAG GTTTTTTTTTTTTVN 3' |
| Poly(T) adaptor specific reverse primer | 5' GCGAGCACAGAATTAATACGAC 3' |
| U6 Forward primer | 5' GCTCGCTTCGGCAGCACA 3' |
| U6 Reverse primer | 5' AACGCTTCACGAATTTGCGTG 3' |
| ID1 3'-UTR Wt sense oligo: | 5' CTAGACGCTGAAGCGCCTCCCCCAGGGACCG GCGGACCCCAGCCATCCAGGGGGCAAGAGG AATTACGTGCTC 3' |
| ID1 3'-UTR Wt antisense oligo: | 5' CTAGAGAGCACGTAATTCCTCTTGCCCCCTG GATGGCTGGGGTCCGCCGGTCCCTGGGGGAG GCGCTTCAGCG 3' |
| ID1 3'-UTR Mut sense oligo: | 5' CTAGACGCTGAAGCGCCTCCCCCAGGGACCG GCGGAGGGGAGCCATCCAGGGGGCAAGAGG AATTACGTGCTC 3' |
| ID1 3'-UTR Mut antisense oligo: | 5' CTAGAGAGCACGTAATTCCTCTTGCCCCCTG GATGGCTCCCCTCCGCCGGTCCCTGGGGGAG GCGCTTCAGCG 3' |
| ID1 3'-UTR RIP Forward | 5' AGAGGAATTACGTGCTCTGTG 3' |
| IDI 3' UTR RIP Reverse | 5' AGGCTGGATGCAGTTAAGG 3' |
